# Head-to-head comparison of nuclear imaging approaches to quantify tumor CD8^+^ T-cell infiltration

**DOI:** 10.1101/2024.10.13.618082

**Authors:** Gerwin G.W. Sandker, René Raavé, Inês F. Antunes, Milou Boswinkel, Lenneke Cornelissen, Gerben M. Franssen, Janneke Molkenboer-Kuenen, Peter J. Wierstra, Iris M. Hagemans, Erik F.J. de Vries, Johan Bussink, Gosse Adema, Martijn Verdoes, Erik H.J.G. Aarntzen, Sandra Heskamp

## Abstract

Many immunotherapies focus on (re)invigorating CD8^+^ T cell anti-cancer responses and different nuclear imaging techniques have been developed to measure CD8^+^ T cell distributions. *In vivo* labeling approaches using radiotracers primarily show CD8^+^ T cell distributions, while *ex vivo* labeled CD8^+^ T cells can show CD8^+^ T cell migration patterns, homing, and tumor infiltration. Currently, a comprehensive head-to-head comparison of *in vivo* and *ex-vivo* cell labeling with respect to their tumor and normal tissue targeting properties and correlation to the presence of CD8^+^ T cells is lacking, yet essential for correct interpretation of clinical CD8^+^ imaging applications. Therefore, we performed a head-to-head comparison of three different CD8^+^ T cell imaging approaches: 1) ^89^Zr-labeled DFO-conjugated Fc-silent anti-CD8 antibody ([^89^Zr]Zr-anti-CD8-IgG2a_silent_), 2) *ex vivo* ^89^Zr-oxine labeled ovalbumin-specific CD8^+^ T cells ([^89^Zr]Zr-OT-I cells), and 3) ^18^F-labeled IL2 ([^18^F]AlF-RESCA-IL2).

**Methods:** B16F10/OVA tumor-bearing C57BL/6 mice (n=10/group) received intravenously one of the three radiopharmaceuticals. PET/CT images were acquired starting 72 h ([^89^Zr]Zr-anti-CD8-IgG2a_silent_), 24 and 48 h ([^89^Zr]Zr-OT-I cells), and 10 min ([^18^F]AlF-RESCA-IL2) post injection. Subsequently, *ex vivo* biodistribution analysis of the radiopharmaceuticals was performed followed by flow cytometric analysis to evaluate the number of intratumoral CD8^+^ T cells. Additionally, the intratumoral radiolabel distributions was assessed by autoradiography and immunohistochemistry (IHC) on tumor slices.

**Results:** [^89^Zr]Zr-anti-CD8-IgG2a_silent,_ [^89^Zr]Zr-OT-I cells, and [^18^F]AlF-RESCA-IL2 showed uptake in CD8-rich tissues, with preferential targeting to the spleen. Biodistribution analysis showed tumor uptake above blood level for all radiopharmaceuticals, except [^18^F]AlF-RESCA-IL2. For all three approaches, the uptake in the tumor-draining lymph node was significantly higher compared with the contralateral axial lymph node, suggesting that all approaches allow evaluation of immune responses involving CD8^+^ T cells. Tumor uptake of [^89^Zr]Zr-anti-CD8-IgG2a_silent_ (R^2^=0.65, p<0.01) and [^89^Zr]Zr-OT-I cells (R^2^=0.74, p<0.01) correlated to the number of intratumoral CD8^+^ T cells (flow cytometry). The intratumoral distribution pattern of the radiosignal was different for *ex vivo* and *in vivo* radiolabeling techniques. The short half-life of ^18^F precluded autoradiography assessment of [^18^F]AlF-RESCA-IL2.

**Conclusion:** We show that [^89^Zr]Zr-anti-CD8-IgG2a_silent_ and [^89^Zr]Zr-OT-I cells PET/CT imaging can be used to evaluate intratumoral CD8^+^ T cells, even though their normal tissues and intratumoral distribution patterns are significantly different. Based on their characteristics, [^89^Zr]Zr-anti-CD8-IgG2a_silent_ might be most useful to immunophenotyping the TME, while the *ex vivo* cell labeling approach visualizes CD8^+^ T cell migrations patterns and the permissiveness of tumors for invasion, whereas [^18^F]AlF-RESCA-IL2 allows for rapid recurrent imaging and might prove useful for tracking rapid changes in CD8^+^ T cell distributions. In conclusion, our head-to-head comparison of the three prototype CD8^+^ T cell labeling approaches provides new insights which can aid in correct interpretation of clinical CD8 imaging and may guide in the selection of the optimal imaging approach for the research question of interest.

## Introduction

CD8^+^ T cells are the main effector cells of the adaptive immune system. Therefore, immune therapies often aim to (re)invigorate their anti-cancer responses [1, 2]. It has been shown that the number and the (intratumoral) distribution of CD8^+^ T cells, as measured immunohistochemically, are an important prognostic and predictive factors for treatment success, and thus an important biomarker [1, 3].

CD8^+^ T cells are highly mobile, constantly circulate via lymphatic and vascular systems, and temporarily reside in primary and secondary lymphoid tissues, as well as in non-lymphoid tissues [4, 5]. Previous studies with radiolabeled lymphocytes have shown that under steady state in healthy individuals, T cells mainly reside in the spleen, lymph nodes (LN), and non-lymphoid tissues [6, 7]. Moreover, the distribution of CD8^+^ T cells is determined by their developmental stage as this defines their migration patterns. Naïve CD8^+^ T cells primarily reside in the blood and LNs searching for antigen presenting cells that contain their T cell receptors’ cognate antigen [8], whereas memory T cells remain in non-lymphoid tissues, blood, LNs, or the spleen depending on phenotype [9]. During disease progression (e.g., cancer) the migration of naïve CD8^+^ T cells is halted in the LN by chemotactic and haptotactic factors to facilitate their activation [10]. Upon activation through antigen recognition or immunotherapeutic intervention, several changes occur (e.g., production of interleukin-2 (IL2)) that facilitate anti-cancer effector CD8^+^ T-cell responses. Namely, expansion of the antigen-specific CD8^+^ T-cell clones in the LNs and re-distribution across different tissues including the tumor [2]. Indeed, multiple studies have shown that CD8^+^ T-cell related parameters in the locoregional tumor area are associated with clinical efficacy of immunotherapies. These parameters include increased number of infiltrating cells, location of cells (e.g. core versus rim of the tumor), presence of tertiary lymphoid structures, and expansion of CD8^+^ T-cell population in tumor draining lymph nodes (TDLN) [1, 7, 11, 12]. Moreover, considering that the distribution of CD8^+^ T cells is influenced by their developmental stage, distribution parameters in healthy but non-lymphoid compartments could also provide insights into their functional status. Thus, there is a need for tools to assess *in vivo* distribution parameters of CD8^+^ T cells, as these can inform on possible treatment success for immunotherapies.

Nuclear imaging is uniquely suitable to measure these parameters, as it can measure CD8^+^ T cell distributions longitudinally, quantitatively, and non-invasively on a whole-body scale [13]. Therefore, nuclear imaging can potentially be used as a direct read-out of the aforementioned factors associated with immunotherapy success [7]. Several approaches for imaging CD8^+^ T cells have been developed which can be categorized as 1) T cell marker targeting proteins (e.g. antibodies, minibodies), 2) *ex vivo* labeled T cells, and 3) T-cell targeting cytokines (e.g. IL2) [7, 14–17]. These three prototype imaging approaches inform on distinct aspects of tumor T cell infiltration, focusing on either T cell distributions or T cell migration. A thorough understanding of the tracers’ distribution/kinetics, quantification techniques for PET features, and relations to CD8^+^ T cells in tissues are required to support clinical implementation and interpretation of the data.

Here, in a head-to-head comparison, we investigate these aspects for three distinct imaging approaches to assess tumor infiltration by CD8^+^ T cells, 1) [^89^Zr]Zr-anti-CD8-IgG2a_silent_ antibodies, 2) *ex vivo* [^89^Zr]Zr-OT-I CD8^+^ T cells, and 3) [^18^F]AlF-RESCA-IL2. We performed PET/CT imaging in immunocompetent B16F10/OVA tumor-bearing mice and show that, apart from high spleen uptake, each approach shows a different distribution profile. Furthermore, we observed that tumor uptake of [^89^Zr]Zr-anti-CD8-IgG2a_silent_ and [^89^Zr]Zr-OT-I CD8^+^ T cells correlated with the number of intratumoral CD8^+^ T cells, while their spatial intratumoral distribution shows distinct patterns.

## Methods

### Animals

Female C57BL/6 OT-I/CD90.1^+^ (n=3, 7-8 weeks) mice were bred and housed and female C57BL/6J mice (n=36, 13 weeks, Janvier) were purchased and housed at the Animal Research Facility at the Radboudumc Nijmegen under specific-pathogen-free conditions with cage enrichment present in individually ventilated cages with a filter top (Green line IVC, Tecniplast) with food and water available ad libitum. All *in vivo* experiments were approved by the Central Authority for Scientific Procedures on Animals (AVD1030020173885) and the Animal Welfare Body of the Radboud University, Nijmegen, and were performed following the principles laid out by the Dutch Act on Animal Experiments (2014). Blinding for group allocation was applied for biotechnicians during experimental execution and for researchers until tracer injection. For a graphical overview of the study see supplemental figure 1.

### Tumor cell inoculation

Ovalbumin-expressing B16F10 melanoma tumor cells (B16F10/OVA) were cultured in RPMI-1640 (L0501, Biowest) supplemented with L-glutamine (Gibco), 10% fetal calf serum (FCS, Sigma-Aldrich Chemie BV), and Geneticin sulphate solution (g418, final concentration 1 mg/mL) for positive selection of transfectants. Cells were cultured in a humidified atmosphere with 5% CO_2_ at 37 °C and split when confluency was 80% at maximum. Cells were collected by trypsinization, washed with supplemented culturing medium, and resuspended in 4 °C RPMI-1640. Next, ice cold Matrigel high concentration (354248, Corning) was added to reach a final 1:3 Matrigel:RPMI-1640 solution of 2.5×10^6^ cells/mL. Mice were anesthetized and inoculated subcutaneously on the right shoulder with 100 µL RPMI-1640/Matrigel containing 0.25×10^6^ B16F10/OVA cells. Tumor size was measured twice weekly using a caliper. Prior to injection, 9 days post inoculation, mice with tumor sizes closest to the mean were block randomized based on tumor size (mean size 80.6±19.4 mm^3^) in three groups (n=10/group).

### ^89^Zr-labeling of anti-CD8-IgG2a_silent_

Anti-CD8-IgG2a_silent_, an engineered chimeric antibody with LALA edited mouse IgG2a for silencing of Fc-effector functions, was developed and DFO-conjugated in house and was shown to specifically target CD8^+^ T cells *in vivo* without target-cell depletion (data will be published elsewhere). First, pH of [^89^Zr]Zr oxalate in 1M oxalic acid (61.2 MBq, NEZ308000MC, PerkinElmer) was adjusted by adding Na_2_CO_3_ to a final concentration of 0.62 M and incubated at room temperature (RT) for 3 min. Next, we added HEPES pH 7.5 to a final concentration of 0.39 M. Subsequently, 104 µg DFO-anti-CD8-IgG2a_silent_ was added and incubated under metal free conditions for 40 min at 300 rpm and 37 °C on a thermoshaker. Labeling efficiency (LE) was determined with instant thin-layer chromatography (iTLC), using silica gel chromatography strips (Agilent Technologies) and 0.1 M, pH 6.0 citrate (Sigma-Aldrich) as running buffer and evaluated using photosensitive plates (Fuji multipurpose standard (MS), Cytiva), a phosphor imager (Typhoon FLA 7000, GE), and AIDA v.4.21.033 software (Raytest, Straubenhardt). LE of [^89^Zr]Zr-anti-CD8-IgG2a_silent_ was 95.3% at a specific activity of 0.47 MBq/µg. The injection solution of 42.5 µg/mL [^89^Zr]Zr-anti-CD8-IgG2a_silent_ was prepared by adding sterile pyrogen-free phosphate buffered saline (PBS). Mice (n=10, nine days after B16F10/OVA inoculation) were injected intravenously (iv) with 8.5 µg (4.1 MBq) [^89^Zr]Zr-anti-CD8-IgG2a_silent_.

### Isolation and ^89^Zr-labeling of CD8^+^ OT-I T cells

C57Bl/6 OT-I/CD90.1^+^ donor mice were sacrificed by cervical decapitation. Spleen and inguinal lymph nodes were transferred in PBS at 4 °C and prepared into single cell suspensions using cell strainers (100 µm) and PBS supplemented with 2% fetal calf serum (FCS) and 1 mM ethylenediaminetetraacetic acid (EDTA) (PFE). Spleen and lymph node cells were pooled, spun down, and resuspended at 1×10^8^ cells/mL in PFE. MHC class I-restricted, ovalbumin-specific, CD8^+^ T cells (OT-I cells) were isolated by negative selection with the EasySep^TM^ Mouse CD8 T cell isolation kit (Catalog #19853, Stemcell Technologies) following the manufacturers protocol. Flow cytometry showed an isolation purity of 93% OT-I cells whereas trypan blue staining showed a viability of 95.1% [18].

OT-I cells were labeled using passive membrane diffusing [^89^Zr]Zr(8-hydroxyquinoline)_4_ by the one-step procedure developed by Man *et al* [19]. For this procedure the labeling formulation used was a sterile-filtered solution consisting of 0.5 mg/mL 8-hydroxyquinoline, 1 M HEPES, and 1 mg/mL polysorbate-80 in H_2_O (10 M NaOH was used to set pH to 7.9–8.0). To 100 µL of this solution, 18 µL [^89^Zr]Zr oxalate in 1M oxalic acid (21.8 MBq) was added, briefly mixed, and incubated for 5 min at RT. Labeling efficiency was determined by ITLC using Whatman no.1 strips as solid phase and ethyl acetate (109623, Merck) as running buffer. The labeling efficiency of ^89^Zr-oxine was 72.6%.

The isolated OT-I T cells were spun down (5 min, 270 g) and resuspended in PBS to a concentration of 14.8×10^6^ cells/mL. Next, ^89^Zr-oxine (8 MBq) was added in a 1:30 ratio of the cell suspension volume and incubated for 15 min at 37 °C on a thermomixer. After incubation, PBS was added to a reach a final volume of 5 mL followed by gently resuspending the cells. Unincorporated ^89^Zr was removed by centrifugation for 5 min at 270g and resuspension of the cell pellet in 5 mL fresh PBS. This was repeated (three times) until the activity in the supernatant of the purified sample was reduced to a maximum of 10%. The yield was 0.08 MBq/10^6^ cells. Prior to injection, cells were resuspended in PBS to a final concentration of 9.9×10^6^ cells per mL. Cell viability was determined shortly before injection using trypan blue and indicated 89% viability. Mice (n=10, 10 days after B16F10/OVA inoculation) were injected iv with 2×10^6^ [^89^Zr]Zr-OT-I cells (0.16 MBq).

### ^18^F-labeling of IL2

The fluorine-18 radiolabeling of IL2 was performed as described by van der Veen *et al* [20] using the indirect AluminumFluoride-18-RESCA ([^18^F]AlF-RESCA, [21]) methodology. Briefly, [^18^F]AlF (4.25 GBq) was allowed to react with the RESCA-tetrafluorophenol ester ((6)- H_3_RESCA-TFP, Leuven University) for 15 min at RT. The LE was evaluated with ITLC as described above except using 75% ACN as running buffer, and was 17.7%. Next, [^18^F]AlF-RESCA-TFP (3.18 GBq) was purified using an HLB cartridge (WAT094225, Oasis) and eluted with 0.6 mL ethanol followed by 0.7 mL sodium acetate (pH 8.5). Then, 200 µg IL2 in 100 µL DMSO was added to the eluted [^18^F]AlF-RESCA-TFP (291 MBq) and was conjugated by incubation for 15 min at 50 °C. The conjugate was purified using a tC2 plus cartridge and eluted using 0.6 mL ethanol containing 0.25% H_3_PO_4_ and 5 mL of 5% glucose solution containing 0.1% sodium dodecyl sulfate, resulting in 15.37 MBq of [^18^F]F-RESCA-IL2 ([^18^F]F-IL2). The purity was determined with ITLC as described above except using 0.9% NaCl as running buffer, and was 99.2%. 12 days after tumor cell inoculation, and 10 min before PET/CT imaging, tumor-bearing mice (n=10) were injected iv with 200 µL of eluate containing equal amounts of IL2 protein (7.1 µg (assuming no loss during filtration)) and, depending on time of injection, a radiation dose of 0.473 to 0.163 MBq [^18^F]AlF-RESCA-IL2.

### PET/CT imaging

Because of the different pharmacokinetics of each radiotracer, PET scans were acquired (60 minutes acquisition time) at previously determined (optimal) time points for each specific radiotracer [16, 20]: starting at 72 h pi for [^89^Zr]Zr-anti-CD8-IgG2a_silent_, 24 and 48 h pi for [^89^Zr]Zr-OT-I cells, and at 10 min for [^18^F]F-IL2. Mice were scanned under inhalation anesthesia (2% isoflurane in oxygenated air) and body temperature was kept at 38 °C. Up to four mice, each tape-fixated on a Perspex plate, were scanned simultaneously. To enable quantification, a 5 min transmission scan using a cobalt-57 source was performed for attenuation correction. Immediately following PET scanning using an Inveon animal PET scanner (Siemens Preclinical solutions), while still fixated on Perspex plates, each animal was transferred individually to a SPECT/CT scanner (U-SPECT II/CT, MILabs) for subsequent anatomical CT imaging with 160 µm spatial resolution, 615 µA, and 65 kV as parameters. PET data was reconstructed using Inveon Acquisition Workspace software version 1.5 (Siemens Preclinical Solution) with the reconstruction algorithm OSEM3D/SO-MAP and the following settings: no scatter correction; using scale factor, 0; matrix, 256 × 256; image zoom, 1; frames, all or 4 ([^18^F]AlF-RESCA-IL2); OSEM Iterations, 2; MAP Iterations, 18; target resolution, 1.5 mm; voxel size, 0.4×0.4×0.8 mm. CT data were reconstructed using the MILabs software version 2.04. For subsequent analyses we used the full time window for [^89^Zr]Zr-anti-CD8-IgG2a_silent_ and [^89^Zr]Zr-OT-I cells and the 45 to 60 minutes time window for [^18^F]AlF-RESCA-IL2. The PET and CT images were merged, and Inveon Research Workplace software version 4.1 (Siemens) was used to create maximum-intensity projections (MIP). For quantification of the tumor uptake, volumes of interest (VOIs) enveloping the tumor tissue were drawn based on the CT signal. The adjacent tumor draining lymph nodes were excluded based on visual interpretation of both CT and PET images. Next, threshold limits of activity (in Bq/mL) for inclusion were based on the decay corrected highest voxel intensity observed within each group. The upper thresholds were set at 100% voxel intensity while the lower thresholds were determined at 5% and 40% (for A_5_ and A_40_, respectively), respectively. A_max_ was based on the highest voxel intensity within the VOI. Then, standardized uptake values (SUVs) were calculated by correcting A_5_, A_40_, or A_max_ for the injected dose and body weight of each animal. One SUV_40_ data point of OT-I T cells was excluded as its small VOI size prohibited its correct calculation.

### Biodistribution analysis

To quantify the tissue uptake of [^89^Zr]Zr-anti-CD8-IgG2a_silent_, [^89^Zr]Zr-OT-I cells, and [^18^F]F-IL2 we performed *ex vivo* biodistribution analyses. After PET/CT scanning, mice were euthanized by CO_2_/O_2_ asphyxiation, immediately followed by heart puncture to collect blood and extraction of spleen and tumor tissues. Tumors were halved. One tumor half and the spleen were embedded in Tissue-Tek (4583, Sakura) and snap frozen in isopentane (−80 °C) for immunohistochemical and autoradiographic analyses. The other tumor half was transferred to ice cold PBS for subsequent flow cytometric analysis. Next, other relevant tissues were collected (muscle, lung, thymus, kidney, liver, duodenum, colon, brown adipose tissue, femur, bone marrow, knee, axillary lymph node, inguinal lymph node, and tumor draining lymph node). Blood and tissues samples were weighed, and radioactivity concentrations were measured with a well-type gamma-counter (Wallac 2480 wizard, Perkin Elmer) or a NaI crystal scintillation counter (Osprey, Mirion) for tumor and spleen tissues. For reference, we used concomitantly measured aliquots of injection fluid or a calibration curve in case of [^89^Zr]Zr-anti-CD8-IgG2a_silent_. Tissue accumulation of [^89^Zr]Zr-anti-CD8-IgG2a_silent_, [^89^Zr]Zr-OT-I cells, and [^18^F]F-IL2 was calculated as a percentage of the injected dose and normalized for tissue weight (percentage injected dose per gram tissue (%ID/g)) and SUV (see supplemental table 2).

### Flow cytometry for intratumoral CD8^+^ T cells

We investigated the number intratumoral CD8^+^ T cells with flow cytometry using the T cell markers CD45 and CD8. Tumor tissue was converted to single cell suspensions using cell strainers (70 µm) and PBS containing 2% FCS and 2 mM EDTA (E-1644, Sigma). Cells were centrifuged at 515g for 5 min, washed with PBS, and resuspended in PBS containing 1% FCS and 2 mM EDTA. In duplicate 1×10^6^ cells were transferred to 96 wells plates, centrifuged at 400g for 3 min, incubated with Efluor780 (Fixable Viability Dye eFluor 780, 65-0865-18, eBioscience) diluted 1:1000 in 100 µL PBS for 10 min at RT. Next, cells were washed and incubated with 100 µL PBS containing 1% FCS, 2mM EDTA, 0.833 µg/mL rat-anti-mouse-CD45-FITC (clone 30-F11, 103108, Biolegend) and 0.333 µg/mL rat-anti-mouse CD8alpha-APC (clone 53-6.7, 100711, Biolegend) for 30 min at 4 °C. Control samples were concomitantly incubated with PBS containing 1% FCS, 2mM EDTA containing no marker, or one of the antibodies or eFluor780. Thereafter, cells were washed three times, resuspended in PBS containing 1% FCS, 2mM EDTA and analyzed with a flow cytometer (FACSLyric, BD Bioscience). Gating was performed for viable cells, CD45^+^, and CD8^+^ for selection of CD8^+^ lymphocytes (supplemental figure 2).

### Immunohistochemistry

Frozen tumor sections (5 µm, n=5/group) were evaluated by immunohistochemistry for CD8, F4/80, and blood vessels (marker 9F1). Sections were fixed in acetone (−20 °C) for 10 min, air dried, and subsequently washed in PBS. For CD8, primary antibody incubation was performed using rat-anti-mouse CD8alpha (KT15, MAS-16761, Thermo Scientific) diluted 1:200 in primary antibody diluent (PAD, 926001, Biolegend) for 2 h at RT. Next, we blocked rat-epitopes with goat-anti-rat-FAB (112-007-003, Jackson ImmunoResearch) at 1:100 dilution in PAD for 45 min at RT. Subsequently, secondary antibody donkey-anti-goat-Alexa488 (A11055, Invitrogen) diluted 1:600 in PAD was incubated for 45 min at RT. Next, for blood vessels staining, sections were incubated with primary antibody rat-anti-mouse-CD146 (9F1) ready-to-use in culturing supernatant for 45 min at 37 °C, followed by secondary antibody chicken-anti-rat-Alexa647 (A21472, Invitrogen) diluted 1:100 in PAD for 45 min at 37 °C. After blocking endogenous biotin (SP-2001, Vector) using the manufacturers protocol, we incubated with rat-anti-mouse biotinylated-F4/80 (13-4801-85, eBioscience) diluted 1:100 in PAD for 45 min at 37 °C and subsequently with secondary antibody mouse-anti-biotin-Cy3 (200-162-211, Jackson ImmunoResearch) diluted 1:100 in PAD for 30 min at RT. Sections were mounted using DAPI-containing fluoromount-G (00-4959-52, Invitrogen) and a cover slip. Sections were scanned at 20x objective magnification acquired using a Zeiss Axioskop fluorescence microscope (Zeiss) using IP-lab imaging software (Scanalytics). Images from representative tumor areas were created using Fiji version 1.53t.

### Autoradiography

Frozen tumor sections (5 µm, consecutive to IHC) of [^89^Zr]Zr-anti-CD8-IgG2a_silent_ and [^89^Zr]Zr-OT-I cells injected mice were exposed to photosensitive plates (BAS-IP super resolution, Fuji, Cytiva). These were scanned using a phosphor imager (Typhoon FLA 7000, GE) at a size of 25×25 pixels/mm and analyzed using AIDA v.4.21.033 software (Raytest). General co-localization of the radiosignal with CD8^+^ cells, F4/80^+^ cells, and vessels was performed by visual evaluation. The short half-life of ^18^F prohibited autoradiographic distribution analysis of [^18^F]AlF-RESCA-IL2.

### Statistical analysis

Data from previous experiments (unpublished data) was used to determine the required number of animals with *a priori* sample size calculations. Differences in tissue uptake between the approaches and differences in ratios was statistically tested for significance using one-way or two-way ANOVAs with Tukey’s correction or one-way ANOVAs with Holm-Šídák correction for multiple testing, respectively. To test if ratios significantly deviated from one, 95% confidence intervals (95%CI) were calculated. Data is shown as mean±standard deviation (SD) unless stated otherwise. Statistical significance was defined as p<0.05. Relation between tumor uptake and abundance of CD8^+^ T cells was evaluated with Pearson correlation coefficients. Correlations were defined as no (R^2^: <0.3), moderate (R^2^: 0.3-0.5), or high (R^2^: 0.5-1) correlation provided that p<0.05.

## Results

### General biodistributions

The biodistribution of [^89^Zr]Zr-anti-CD8-IgG2a_silent_, [^89^Zr]Zr-OT-I CD8^+^ T cells, and [^18^F]AlF- RESCA-IL2 was evaluated in B16F10/OVA tumor-bearing mice under the same experimental conditions using PET/CT imaging at different time points (72 h, 24/48 h, and 55–70 min time range pi, respectively), followed by ex vivo biodistribution studies. PET/CT imaging showed clear tumor uptake for [^89^Zr]Zr-anti-CD8-IgG2a_silent_, whereas tumor uptake of [^89^Zr]Zr-OT-I cells and [^18^F]F-IL2 was difficult to discern above background. High uptake of [^89^Zr]Zr-anti-CD8-IgG2a_silent_ and [^89^Zr]Zr-OT-I cells was observed in CD8-rich tissues such as the spleen, lymph nodes, and duodenum (figure 1), with moderate uptake in the liver. Uptake of [^18^F]AlF-RESCA-IL2 was visualized in the spleen, kidney, bladder, liver, and lung.

**Figure 1.**
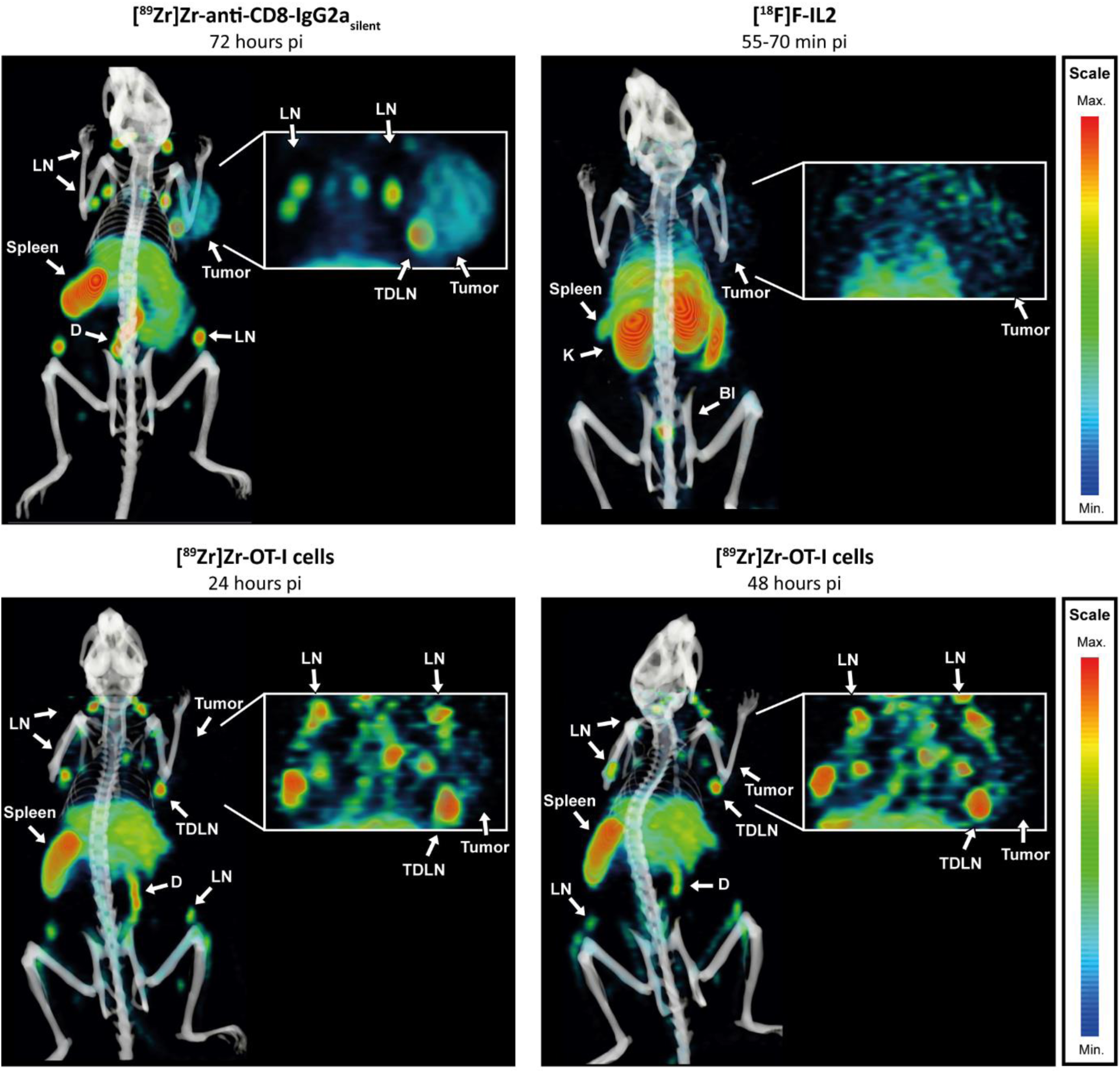
*In vivo* distribution of [^89^Zr]Zr-anti-CD8-IgG2a_silent_, [^89^Zr]Zr-OT-I CD8^+^ T cells, and [^18^F]AlF-RESCA-IL2. PET/CT derived representative MIPs (optimally scaled per approach) visualizing the distribution of 8.5 µg [^89^Zr]Zr-anti-CD8-IgG2a_silent_(72 h pi), 2×10^6^ [^89^Zr]Zr-OT-I cells (24 and 48 h pi), and 0.364 MBq [^18^F]AlF-RESCA-IL2 (55–70 min pi time range) in B16F10/OVA tumor-bearing female C57BL/6J mice. Tissues indicated are tumor, spleen, lymph nodes (LN), tumor draining lymph node (TDLN), duodenum (D), bladder (Bl) and kidney (K). Inserts have increased intensity scaling to visualize tumor uptake.

*Ex vivo* biodistribution studies confirmed these findings (complete biodistribution data is presented in supplemental table 1). Blood levels were higher for [^18^F]AlF-RESCA-IL2 (2.8±0.7 %ID/g) compared with [^89^Zr]Zr-anti-CD8-IgG2a_silent_ (1.5±0.1 %ID/g) and [^89^Zr]Zr-OT-I T cells (0.4±0.0 %ID/g). The moderate bone uptake of each radionuclide indicated a limited degree of ^89^Zr and ^18^F de-chelation.

In summary, [^89^Zr]Zr-anti-CD8-IgG2a_silent_, [^89^Zr]Zr-OT-I cells, and [^18^F]AlF-RESCA-IL2 all showed moderate tumor uptake whereas the former two also preferentially accumulated in all CD8-rich tissues. High [^18^F]AlF-RESCA-IL2 uptake was observed in the spleen, excretory organs, and lungs.

### Tumor uptake – *ex vivo* biodistribution assessed

To evaluate the feasibility of each approach to evaluate CD8^+^ T cell tumor infiltration, we compared their tumor uptake and tumor-to-background ratios. The mean tumor sizes were equal between the three groups at time of block-randomization, however, a day before dissection tumor sizes of the [^89^Zr]Zr-anti-CD8-IgG2a_silent_ group (241±86 mm^3^, supplemental figure 3) were smaller compared with those of [^89^Zr]Zr-OT-I (415±288 mm^3^, p<0.001) and [^18^F]AlF-RESCA-IL2 (351±159 mm^3^, p=0.037). Tumor uptake for [^89^Zr]Zr-anti-CD8-IgG2a_silent_, [^89^Zr]Zr-OT-I cells and [^18^F]AlF—RESCA-IL2 was 3.4±1.5, 1.9±0.4, and 1.7±0.3 %ID/g, respectively (p=ns; Figure 2A). Tumor-to-muscle ratios were >1 for all tracers and due to its low muscle uptake highest for [^89^Zr]Zr-anti-CD8-IgG2a_silent_ (31.9±16.8, p<0.001 for both comparisons), followed by [^89^Zr]Zr-OT-I cells (3.2±1.6) and [^18^F]AlF-RESCA-IL2 (1.7±0.4; figure 2A). Tumor-to-blood ratios were higher for [^89^Zr]Zr-OT-I cells (4.3±1.2) compared with [^89^Zr]Zr-anti-CD8-IgG2a_silent_ (2.2±1.0, p<0.001), whereas the tumor uptake of [^18^F]AlF-RESCA-IL2 was below blood level, suggesting that specific uptake might be difficult to discern.

**Figure 2.**
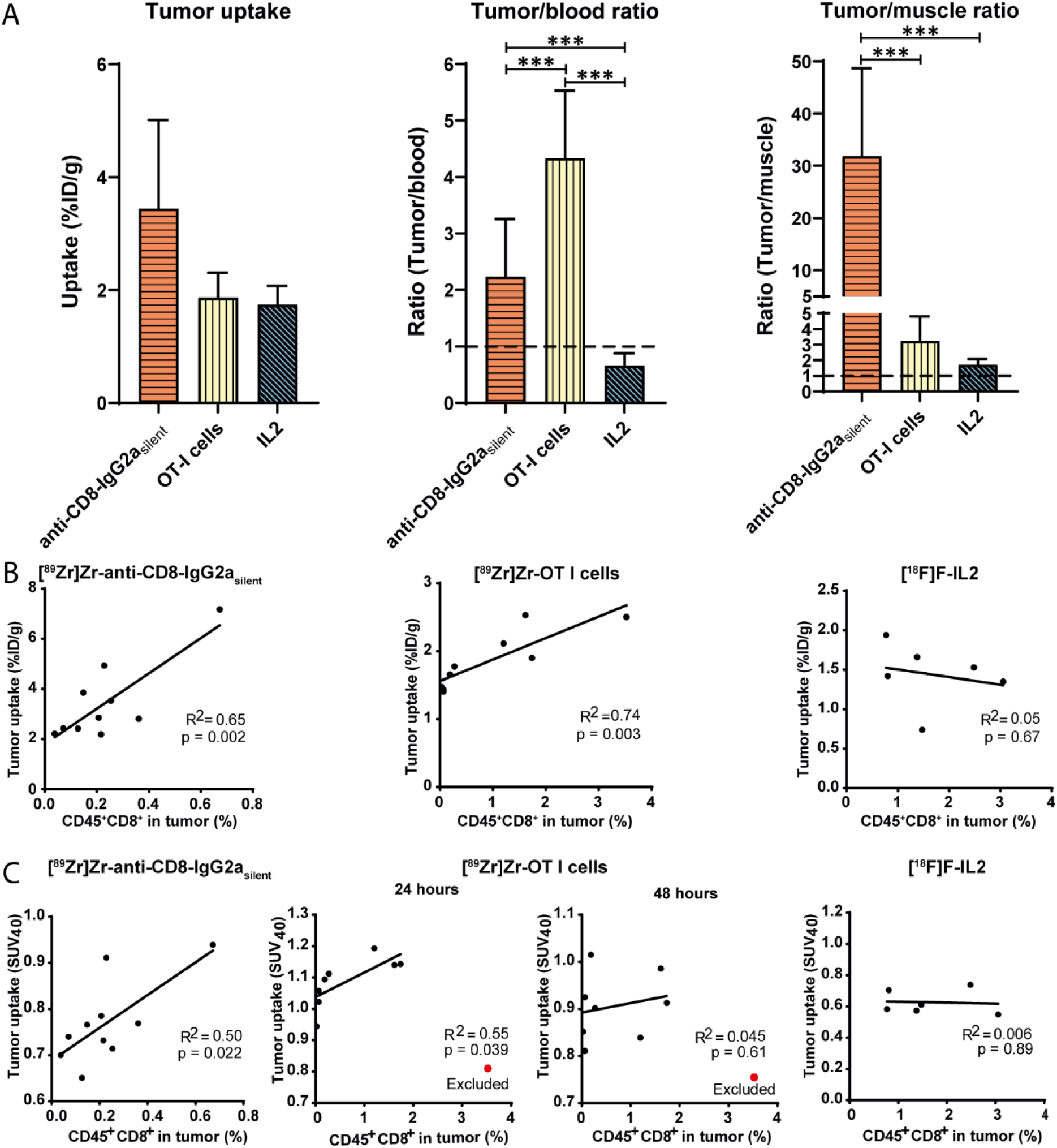
Tumor uptake characteristics and correlations with the number of tumor infiltrated CD8^+^ T cells of [^89^Zr]Zr-anti-CD8-IgG2a_silent_ antibody, [^89^Zr]Zr-OT-I CD8^+^ T cells, and [^18^F]AlF-RESCA-IL2. **(A)** Tumor uptake, Tumor-to-blood, and Tumor-to-muscle ratios of [^89^Zr]Zr-anti-CD8-IgG2a_silent_ antibody (72 h pi, n=10), [^89^Zr]Zr-OT-I cells (48 h pi, n=9), and [^18^F]AlF-RESCA-IL2 (Biodistribution: 75 min pi, PET: 55–70 min pi, n=10) in C57BL/6 mice bearing B16F10/OVA tumors (*** = p<0.001). The correlation of the percentage of CD45^+^CD8^+^ T cells as determined by flow cytometry with their tumor uptake determined by **(B)** *ex vivo* biodistribution analysis and **(C)** PET image quantification. One SUV_40_ data point of OT-I T cells was excluded as a limited VOI size prohibited its correct calculation.

### Tumor uptake – PET quantified

To evaluate the PET quantified tumor uptake of each approach, we assessed their total tumor uptake (SUV_5_), tumor uptake above 40% of the maximum (SUV_40_), and maximum uptake (SUV_max_). SUV_5_ was higher for [^89^Zr]Zr-anti-CD8-IgG2a_silent_ than for [^89^Zr]Zr-OT-I cells, whereas SUV_40_ and SUV_max_ were higher for [^89^Zr]Zr-OT-I cells (table 1 and supplemental table 2 for a detailed overview). This indicates a more heterogenous uptake pattern for [^89^Zr]Zr-OT-I cells. Uptake of [^89^Zr]Zr-OT-I cells was slightly higher at 24 h compared with 48 h pi for all measured SUV parameters. In line with biodistribution data, PET quantified radioactive signal showed lowest SUV_5_, SUV_40_, and SUV_max_ in the tumor for [^18^F]AlF-RESCA-IL2.

**Table 1.**
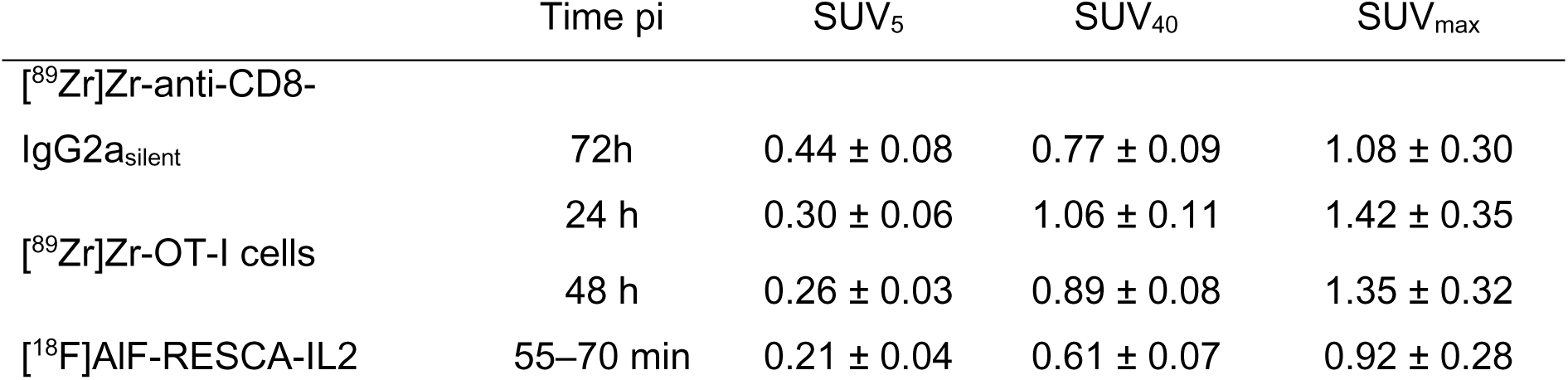
SUV values of tumor uptake. PET quantified SUV_5_, SUV_40_, and SUV_max_ values for tumor uptake of [^89^Zr]Zr-anti-CD8-IgG2a_silent_, [^89^Zr]Zr-OT-I CD8^+^ T cells, and [^18^F]AlF-RESCA-IL2.

### Correlation tumor uptake with number of intratumoral CD8^+^ T cells

For each approach we assessed the relation between tumor uptake and the number of CD8^+^ T cells measured by flow cytometry. The tumor uptake, determined by *ex vivo* biodistribution, of [^89^Zr]Zr-anti-CD8-IgG2a_silent_ antibodies and [^89^Zr]Zr-OT-I cells showed a high correlation with the percentage of CD45^+^CD8^+^ T cells (R^2^=0.65, p=0.002, and R^2^=0.74, p=0.003, figure 2B), while this correlation was absent for [^18^F]AlF-RESCA-IL2 (R^2^=0.05, p=0.67). Uptake of [^89^Zr]Zr-anti-CD8-IgG2a_silent_ antibody and [^89^Zr]Zr-OT-I cells also strongly correlated with the fraction of CD8^+^ cells within the CD45^+^ population (supplemental figure 4).

As determined by PET quantification, the SUV_5_ of [^89^Zr]Zr-anti-CD8-IgG2a_silent_ did not correlate with the percentage of CD8^+^ T cells (R^2^=0.28, p=0.12), while a moderate correlation was observed for [^89^Zr]Zr-OT-I cells at the 24 h (R^2^=0.62, p=0.02) but not the 48 h post injection scan (supplemental figure 5). SUV_5_ values for [^18^F]AlF-RESCA-IL2 did not correlate with the percentage of CD8^+^ T cells in the tumor. In contrast, SUV_40_ showed a high positive correlation with the percentage of CD8^+^ T cells for both [^89^Zr]Zr-anti-CD8-IgG2a_silent_ (R^2^=0.50, p=0.022) and [^89^Zr]Zr-OT-I cells at the 24 h pi (R^2^=0.55, p=0.039), whereas no correlation was observed for [^18^F]AlF-RESCA-IL2 (figure 2C). SUV_max_ showed a positive correlations with the percentage of CD8^+^ T cells for [^89^Zr]Zr-anti-CD8-IgG2a_silent_ (R^2^=0.44, p=0.037) and [^89^Zr]Zr-OT-I cells (R^2^=0.78, p<0.01), and no correlation for [^18^F]AlF-RESCA-IL2.

Flow cytometry revealed that while the percentage of viable cells was similar, the percentage of CD45^+^CD8^+^ cells within the viable cell fraction of the tumor was statistically significantly different between [^89^Zr]Zr-anti-CD8-IgG2a_silent_ and [^18^F]AlF-RESCA-IL2 (0.23±0.17 vs 1.66±0.84, p=0.0096, supplemental figure 6) but not between [^89^Zr]Zr-anti-CD8-IgG2a_silent_ and [^89^Zr]Zr-OT-I cells (0.96±1.12, p=ns). This was unrelated to the tumor volume (see supplemental figure 7).

In summary, tumor uptake of [^89^Zr]Zr-anti-CD8-IgG2a_silent_ and [^89^Zr]Zr-OT-I cells correlated positively with the number of CD8^+^ T cells in the tumor, whereas [^18^F]AlF-RESCA-IL2 uptake did not correlate with the number of CD8^+^ T cells.

### Spatial distribution of the radiosignal and CD8^+^ T cells

We assessed the intratumoral tracer distribution using sagittal PET derived slices, and by comparing the autoradiographic signal with the presence of CD8^+^ T cells as assessed by IHC. Sagittal PET derived tumor slices and autoradiography revealed similar distribution patterns (figure 3A-B). Namely, tumor uptake of [^89^Zr]Zr-anti-CD8-IgG2a_silent_ was heterogenous and primarily located at the tumor rim, whereas [^89^Zr]Zr-OT-I cells tumor uptake was observed throughout the tumor in focal hotspots. For [^18^F]AlF-RESCA-IL2 a heterogenous tumor uptake was observed, primarily located at the tumor rim. Visual inspection of ARG images demonstrated that areas with high [^89^Zr]Zr-anti-CD8-IgG2a_silent_ radioactive signal colocalized with areas with more dense CD8^+^ T cells (Figure 3B), however, this was not a perfect co-localization. The focal uptake of [^89^Zr]Zr-OT-I cells demonstrated by ARG also correlated to denser CD8^+^ T cell areas, although CD8^+^ T cell presence was not limited to areas showing [^89^Zr]Zr-OT-I cell uptake. Additionally, no apparent correlation was observed for both [^89^Zr]Zr-OT-I cell and [^89^Zr]Zr-anti-CD8-IgG2a_silent_ ARG with F4/80 expression or large blood vessels. In summary, [^89^Zr]Zr-anti-CD8-IgG2a_silent_ and [^18^F]AlF-RESCA-IL2 primarily targeted the rim of the tumor, whereas [^89^Zr]Zr-OT-I cells showed focal targeting throughout the tumor. For [^89^Zr]Zr-anti-CD8-IgG2a_silent_ and [^89^Zr]Zr-OT-I cells, tumor uptake partly colocalized with the presence of CD8^+^ T cells.

**Figure 3.**
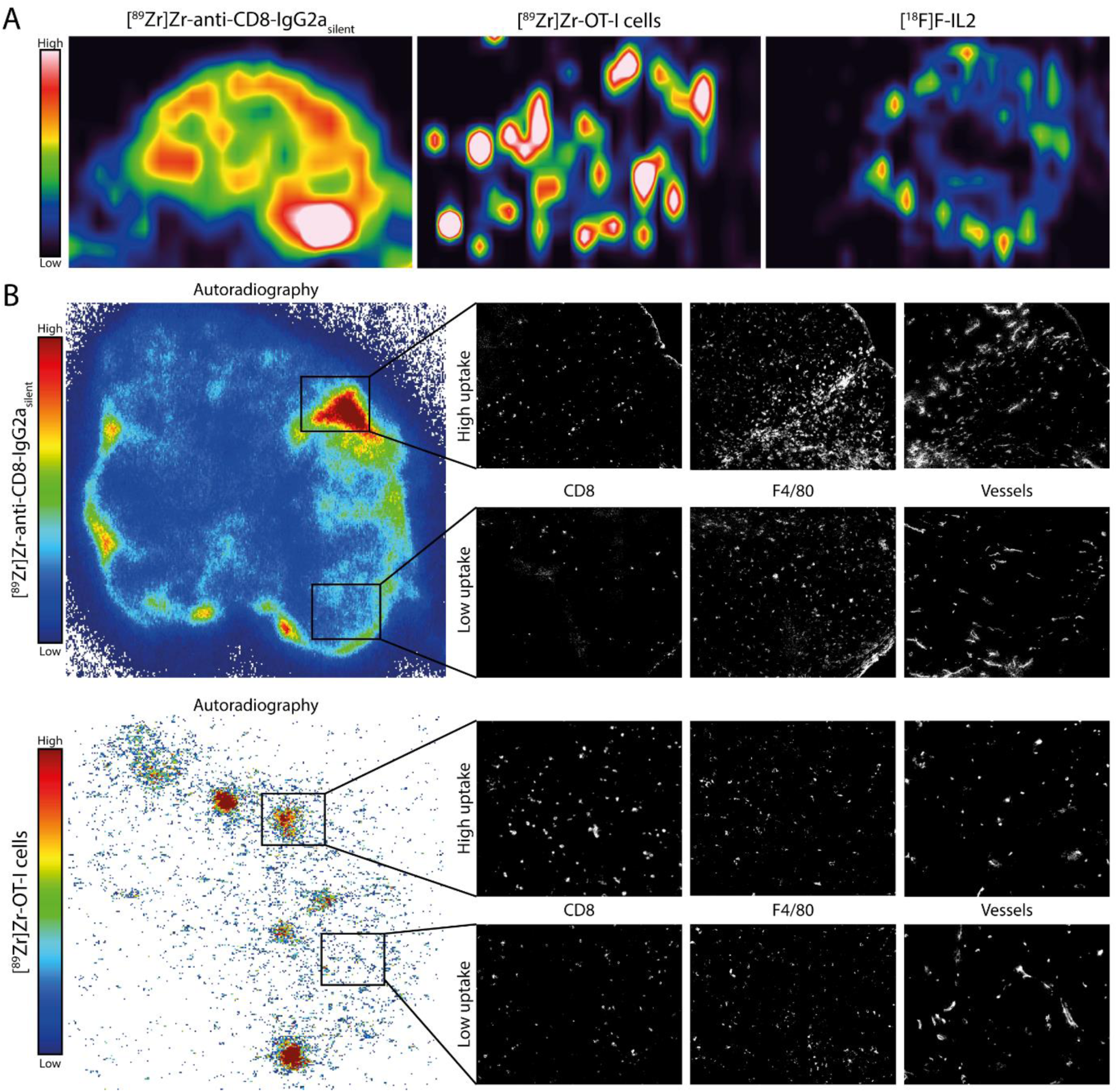
Intratumoral distribution of [^89^Zr]Zr-anti-CD8-IgG2a_silent_, [^89^Zr]Zr-OT-I CD8^+^ T cells, and [^18^F]AlF-RESCA-IL2. Distribution of [^89^Zr]Zr-anti-CD8-IgG2a_silent_ antibody (72 h pi), [^89^Zr]Zr-OT-I cells (48 h pi), and [^18^F]AlF-RESCA-IL2 (55–70 min pi) investigated with PET data derived sagittal tumor sections and *ex vivo* autoradiography compared with the CD8^+^ T cells distribution investigated with IHC on a consecutive section of B16F10/OVA tumors from C57BL/6 mice. **(A)** Representative PET images derived from a sagittal tumor sections. **(B)** Representative images of autoradiography of [^89^Zr]Zr-anti-CD8-IgG2a_silent_ and [^89^Zr]Zr-OT-I T cells (left) and IHC for CD8, F4/80 (center), and blood vessels (right). Of note, the PET images are of a different plane than the autoradiograph and IHC images.

### Uptake in lymphoid compartments

To compare the uptake of the tracers in lymphoid organs, we evaluated their tissue weight corrected uptake, tissue-to-background ratios, and absolute (non-weight corrected) uptake. All radiotracers demonstrated uptake in CD8-rich tissues. For [^89^Zr]Zr-anti-CD8-IgG2a_silent_ and [^89^Zr]Zr-OT-I CD8^+^ T cells, the high splenic (251.1±26.5 and 584.9±53.0 %ID/g, figure 4A) and lymph node uptake (92.3±26.0 and 139.3±30.5 %ID/g) combined with the low concentrations in blood and muscle resulted in high tissue-to-blood and tissue-to-muscle ratios (figure 4B and 4C). For [^18^F]AlF-RESCA-IL2 spleen-to-blood and spleen-to-muscle ratios were above one whereas the low lymph node uptake combined with the relatively high blood concentrations resulted in lymph node-to-blood ratios that did not exceed one. The weight corrected uptake (%ID/g) was higher in the TDLN compared with the contralateral axial lymph node for [^89^Zr]Zr-anti-CD8-IgG2a_silent_ and [^89^Zr]Zr-OT-I cells, but not for [^18^F]AlF-RESCA-IL2. Evaluation of the weight and absolute uptake, expressed as a percentage of the injected dose (%ID), showed higher TDLN weight and uptake when compared with the contralateral LN for all three tracers (figure 4 D). The absolute (%ID) distribution across different tissues within the lymphoid system indicated different distribution patterns except for the predominant accumulation within the spleen for all tracer (figure 5). In summary, for [^89^Zr]Zr-anti-CD8-IgG2a_silent_ and [^89^Zr]Zr-OT-I cells showed high uptake in the spleen and lymph nodes, whereas for each approach higher absolute (but not tissue weight corrected) uptake was observed in the TDLN compared with the contralateral lymph node. Furthermore, within the lymphoid system we observed a different distribution but a preferential localization to the spleen by all tracers.

**Figure 4.**
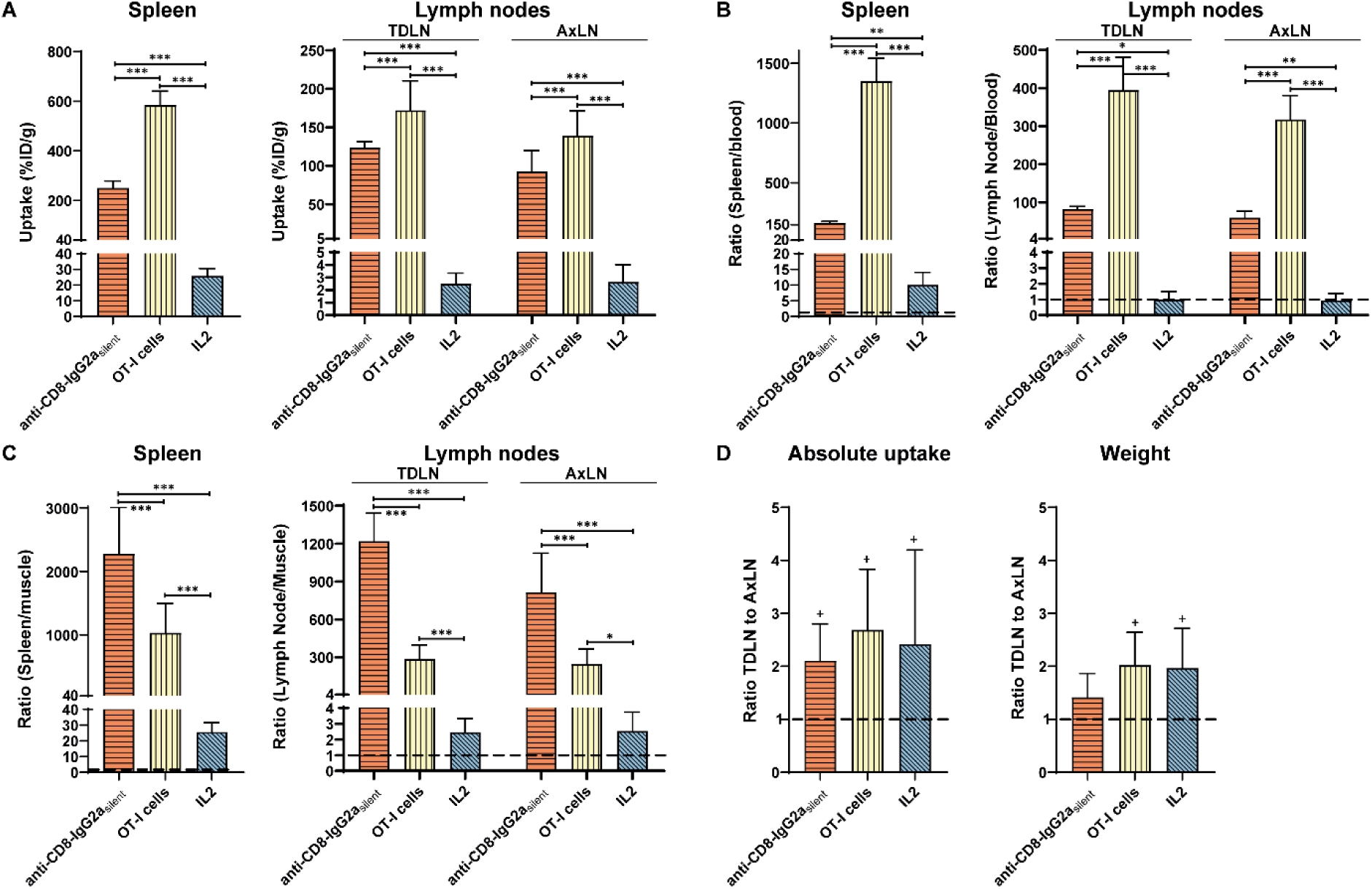
Lymphoid tissue uptake, tissue-to-background ratios, and tumor-draining versus contralateral lymph node uptake ratio. Uptake and uptake ratios of [^89^Zr]Zr-anti-CD8-IgG2a_silent_ (72 h pi, n=10), [^89^Zr]Zr-OT-I CD8^+^ T cells (48 h pi, n=9), and [^18^F]AlF-RESCA-IL2 (75 min pi, n=10) in B16F10/OVA tumor-bearing C57BL/6J mice in the spleen, the tumor draining axial lymph node (TDLN), and its contralateral axial lymph node (AxLN). Shown are the **(A)** tissue uptake, based on tissue weight-normalized data (%ID/g), **(B)** tissue-to-blood ratio, **(C)** the tissue-to-muscle ratio, and **(D)** the ratios between the TDLN and the AxLN, based on the absolute uptake (%ID) and weight. (* = p<0.05, ** = p<0.01, *** = p<0.001, and + = 95%CI does not include 1).

**Figure 5.**
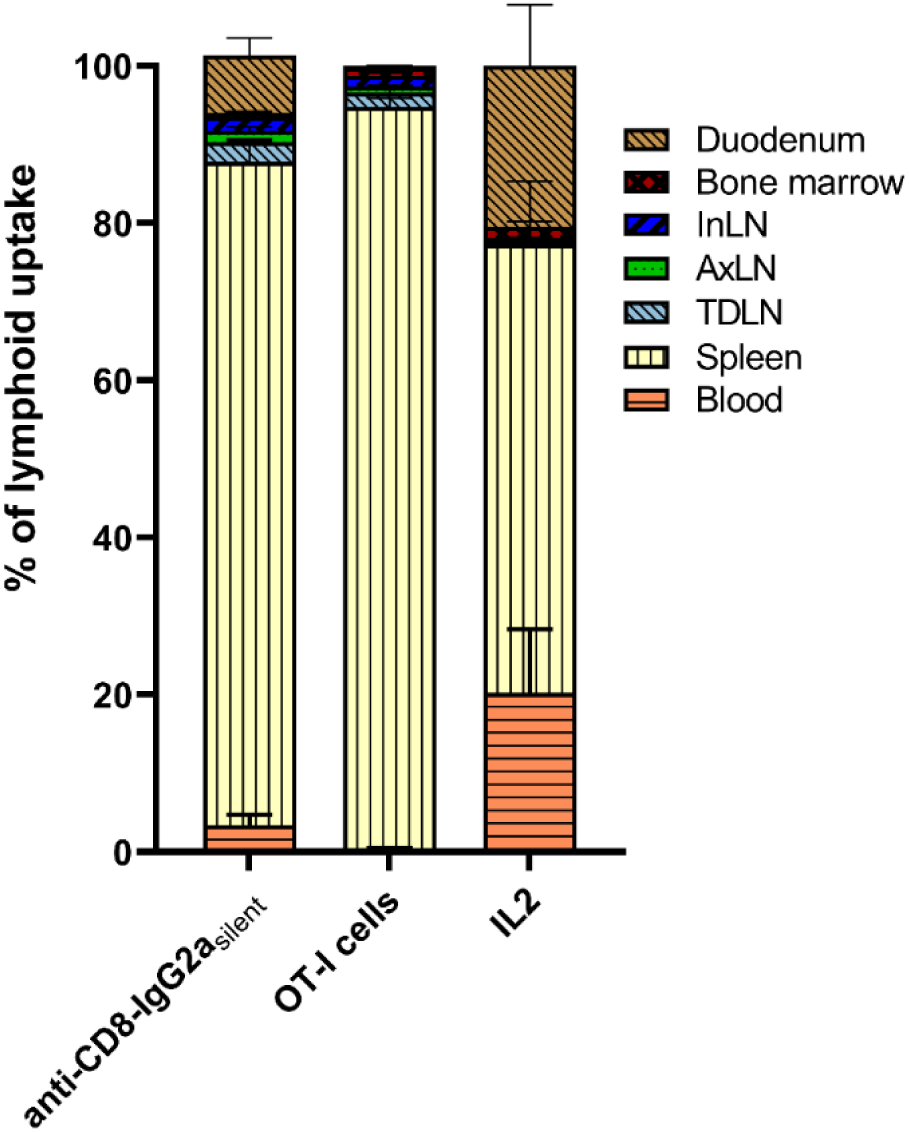
Distribution within lymphoid compartment. Relative lymphoid organ uptake within the total lymphoid compartment of [^89^Zr]Zr-anti-CD8-IgG2a_silent_ (72 h pi, n=10), [^89^Zr]Zr-OT-I CD8^+^ T cells (48 h pi, n=9), and [^18^F]AlF-RESCA-IL2 (75 min pi, n=10) in B16F10/OVA tumor-bearing C57BL/6J mice. Shown are blood, spleen, tumor-draining lymph node (TDLN), contralateral axial lymph node (AxLN), inguinal lymph node (InLN), bone marrow, and duodenum.

## Discussion

Many immunotherapeutic strategies focus on (re)invigorating CD8^+^ T cell anti-cancer responses [2]. Their efficacy is linked to several CD8^+^ T-cell related factors at initiation of therapy [1]. Different nuclear imaging techniques, both *in vivo* and *ex vivo* labeling approaches, have been developed to study CD8^+^ T cells [22]. *In vivo* labeling approaches use radiotracers targeting CD8^+^ T-cell surface markers (e.g., CD8, IL2 receptor) to evaluate the amount and distribution of CD8^+^ T cells within tissues. Approaches using *ex vivo* labeled T cells, in contrast, show migration patterns, homing, and tumor infiltration of these cells. Both approaches are being employed in clinical studies as potential imaging biomarker to predict and monitor response to immunotherapy [7, 23]. A comprehensive understanding of their *in vivo* characteristics, however, is essential for their correct clinical interpretation and application. Here, we performed a head-to-head comparison of the *in vivo* distributions and (PET-derived) tumor uptake values and its relation to the presence of CD8^+^ T cells for three prototype CD8^+^ T cell imaging approaches: ^89^Zr-labeled anti-CD8 IgG2a_silent_ antibodies, *ex vivo* ^89^Zr-labeled OT-I CD8^+^ T cells, and ^18^F-labeled IL2. We demonstrate that tumor uptake of both [^89^Zr]Zr-anti-CD8-IgG2a_silent_ and [^89^Zr]Zr-OT-I correlate to the number of intratumoral CD8^+^ T cells, even though they differ in intratumoral distribution patterns. Furthermore, we show that each approach preferentially targets CD8-rich tissues (most notably the spleen) and might be used to identify lymph nodes with active T cell priming and proliferation.

PET-based assessment of tumor uptake requires sufficient above-background tumor uptake, especially for intratumoral CD8^+^ T cells as the number of intratumoral target cells and consequently the specific signal might be low. High tumor-background ratios were observed for [^89^Zr]Zr-anti-CD8-IgG2a_silent_ and [^89^Zr]Zr-OT-I cells and their uptake values correlated with the numbers and intratumoral distributions of CD8^+^ T cells, both when assessed with *ex vivo* biodistribution analysis and *in vivo* PET quantification. For [^89^Zr]Zr-OT-I cells this suggests that the abundance of CD8^+^ T cells in the tumor appears linked to its permissiveness for infiltration of CD8^+^ T cells. Although [^18^F]AlF-RESCA-IL2 tumor uptake was in the same order of magnitude, no correlation with the number of CD8^+^ T cells was observed. This is possibly explained by the relatively high amount of tracer circulating in the blood at time of dissection which may have resulted in relatively high non-specific tumor uptake masking the specific uptake. Therefore, a longer wash-out period may be preferred to reduce perfusion effects on the uptake. Other possible explanations are associated to characteristics of IL2 receptors, which are also expressed by other lymphocytes in tumors (e.g., regulatory T cells) and whose isoforms have varying IL2 affinities. Upon activation, regulatory T cells retain their high affinity IL2 receptor isoform, whereas CD8^+^ T cells rapidly switch from high to low affinity IL2 receptors [24]. As the used tumor model is highly immunogenic, activation of the intratumoral CD8^+^ T cells may be expected and as a result, intratumoral CD8^+^ T cells may express IL2 receptor isoforms with decreased affinity. This also implies a reliance on a relative overabundance of CD8^+^ T cells for this approach, an experimental parameter which might not have been met in this study. All in all, based on our study in the B16F10/OVA model, both the *in vivo* CD8-targeting based and *ex vivo* cell labeling approaches appear most suitable to evaluate the presence of intratumoral CD8^+^ T cells *in vivo* using PET/CT.

We further showed that PET/CT imaging might be used to evaluate the intratumoral distributions of both [^89^Zr]Zr-anti-CD8-IgG2a_silent_ and [^89^Zr]Zr-OT-I cells, as PET and autoradiography appear to show comparable distribution patterns, although it must be noted that a head-to-head comparison of these techniques is challenging because this would require assessing the exact same slide and is further hindered by differences in resolutions of both techniques. [^89^Zr]Zr-anti-CD8-IgG2a_silent_ preferentially distributed to CD8^+^ T cell denser areas at the rim of the tumor and might therefore be more reflective of the distribution of CD8^+^ T cells in the tumor, indicating that [^89^Zr]Zr-anti-CD8-IgG2a_silent_ with PET/CT may be used to assess intratumoral CD8^+^ T cell distributions. This is in accordance with clinical results with the one-armed CD8-specific antibody [^89^Z]ZED88082A that could discern spatial distributions intratumorally and subsequently phenotype tumors as immune infiltrated or deserts [7]. Tumor uptake of [^89^Zr]Zr-OT-I cells was concentrated in foci which were spatially correlated to the presence of denser CD8^+^ T cell areas using autoradiography and IHC. Denser CD8^+^ T cell areas were, however, also observed outside of the uptake foci. This suggests that [^89^Zr]Zr-OT-I cells preferentially invaded the tumor at similar regions as endogenous CD8^+^ T cells but does not reflect the entire endogenous CD8^+^ T cell distribution. The focal uptake patterns were also observed using PET/CT, suggesting that this approach can visualize CD8^+^ T cell tumor invasion. Thus, the *ex vivo* labeled CD8^+^ T cells approach appears best suited for assessing the tumors’ permissiveness for CD8^+^ T cell infiltration. [^18^F]AlF-RESCA-IL2 PET imaging showed a somewhat similar intratumoral distribution to [^89^Zr]Zr-anti-CD8-IgG2a_silent_, suggesting that in CD8^+^ T cell specific situations and with better to-background ratios it might inform on intratumoral IL2 receptor-positive T cell distributions. However, the low tumor-to-blood ratio hindered correct interpretation of the PET images, indicating that a longer clearance time of the tracer would be required before the start of the PET scan to allow the specific signal to supersede the non-specific signal. The intratumoral distribution of [^18^F]AlF-RESCA-IL2 could not be confirmed by autoradiography because of the short half-life of ^18^F in combination with the low amount of administered activity. For each approach, we further note that the high target mediated and/or non-specific uptake in spleen, bone, kidney, and liver implies that quantitative imaging of CD8^+^ T cells in tumors located in these tissues, or the vicinity of these organs could be difficult.

Immunophenotyping of the tumor microenvironment in may aid in selecting the optimal type of immunotherapy. For example, immune-inflamed tumors primarily may need CD8^+^ T-cell reinvigoration, while immune-excluded tumors may need overcoming immune excluding barriers, and immune desert tumors may benefit most from therapies that prime and activate CD8^+^ T cells [25]. *In vivo* labeling of CD8^+^ T cells with CD8-targeting tracers might be best suited for immunophenotyping the tumor microenvironment as it combines high detection sensitivity and informing intratumoral spatial distributions CD8^+^ T cell. In contrast, through the inherent nature of the approach, PET imaging using *ex vivo* labeled CD8^+^ T cells primarily informs on the permissiveness for tumor infiltration of CD8^+^ T cells and therefore might guide in selecting immunotherapies that either reinvigorate CD8^+^ T cell responses or overcoming hurdles for invasion. Although less successful in this study, cytokine based approaches such as [^18^F]AlF-RESCA-IL2 might, in contrast to CD8-targeting and *ex vivo* cell labeling-based approaches, prove useful to track changes in CD8^+^ T cell distributions over time, as their short circulatory half-life enables repeated imaging. This is more challenging for antibody-based approaches, as their tumor uptake may reflect a combination of CD8^+^ T cells present in the tissue and CD8^+^ T cells that were labeled elsewhere and migrated to tumors over time. This latter compartment is mostly absent for IL2-bases tracers. All in all, for clinical decision making, the IL2-based approach may be more informative for treatment response monitoring than for predictive imaging. A middle ground between these approaches might also be sought in combining the high specific uptake of antibody-based tracers with a shorter circulation time, such as provided by e.g. minibodies [26, 27].

For lymphoid tissues the biodistribution analyses showed high preferential targeting for [^89^Zr]Zr-anti-CD8-IgG2a_silent_ and [^89^Zr]Zr-OT-I T cells, and although still substantial lower for [^18^F]AlF-RESCA-IL2, suggesting target specificity for each approach. These findings are, in general, in line with the biodistributions reported by others, even though the IL2 dose was higher than in a previously reported study [20, 28, 29]. Furthermore, for all three approaches, we clearly observed increased absolute uptake in TDLNs compared with the contralateral LN, which coincided with an enlargement of the LN and thus suggestive of immune response activation. This shows that each approach is suited to assess the activation status of (loco-regional) LNs even though they evaluate different processes, such as locations of CD8^+^ T cell proliferation and expansion ([^89^Zr]Zr-anti-CD8-IgG2a_silent_ and [^18^F]AlF-RESCA-IL2) or migration patterns and therefore the presence of immunological homing and retaining cues for CD8^+^ T cells (*ex vivo* labeled CD8^+^ T cells). For [^18^F]AlF-RESCA-IL2, lymph node uptake did not exceed blood values hampering uptake evaluation using PET, indicating that PET should be acquired at a later timepoint to allow for clearance of the tracer from blood. However, for [^89^Zr]Zr-anti-CD8-IgG2a_silent_ and [^89^Zr]Zr-OT-I cells we observed remarkably high lymph node-to-background ratios suggesting that these approaches are well suited for PET-based evaluation of CD8^+^ T cell presence within these tissues.

Correct clinical application further requires several approach-specific characteristics. CD8-targeting based tracers should not cause depletion of target cells through immune mediated effects, such as antibody dependent cell-mediated cytotoxicity, as this would hinder immunotherapeutic efficacy. [^89^Zr]Zr-anti-CD8-IgG2a_silent_ was “LALA” mutated to minimize binding to Fcγ-receptors making it a non-depleting antibody [30]. Nevertheless, tumors of [^89^Zr]Zr-anti-CD8-IgG2a_silent_ injected mice were smaller and contained a lower percentage of CD8^+^ T cells. The tumor size might have been a statistical type I error, however, we also observed less enlarged TDLNs compared with that of [^18^F]AlF-RESCA-IL2 and [^89^Zr]Zr-OT-I cell treated mice. As ligation of CD8alpha by the used antibody can affect CD8^+^ T cell activation and proliferatrion, biological effects of the antibody affecting CD8^+^ T cell distributions can not be excluded [31, 32]. For *ex vivo* labeled CD8^+^ T cells, viability is an important parameter as they are amongst the most radiation sensitive leukocytes [33]. Adequate viability of the *ex vivo* labeled OT-I cells is suggested by the high uptake in the spleen, although phagocytic uptake of non-viable OT-I cells or uptake of released [^89^Zr]Zr-phosphate metabolites cannot be excluded [34] and long-term effects of radiolabeling on proliferative capacity or effector function have not been investigated. Furthermore, substantial bone uptake of the bone-seeking radiometal suggests that a proportion of the cells have died, as this is likely the primary source of released ^89^Zr [35]. However, the very high lymph node uptake suggests that a large proportion of the radiolabeled OT-I cells retained their migratory capacity and were viable *in vivo* and thus the uptake largely corresponds to migrated CD8^+^ T cells. Additionally, we should note that the OT-I CD8^+^ T cells used are tumor-antigen specific which could have increased its tumor uptake, while in the clinic endogenous CD8^+^ T cells with a multitude of antigen specificity are typically used.

To generalize and translate these findings, several factors should be considered. First, obvious differences between humans and mice, such as relative presence of immune subpopulations, sizes of relevant tissues, and differences in developmental age skewing their maturation states will all result in differences in CD8^+^ T cell distributions and thus pharmacokinetics and distributions of tracer [36, 37]. Second, it becomes increasingly clear that CD8^+^ T cell numbers and distributions show high inter-patient variability, depending for example on age or prior exposures that affect CD8^+^ T-cells [38, 39]. Therefore, a personalized approach in which migration features over time are evaluated might be favored above general descriptive features derived from whole patient populations. Last, other factors should also be taken into consideration when using *ex vivo* labeled approach. For instance, the tumor specificity and maturation and exhaustion state of the cells, pre-infusion work up of the cells (e.g., stimulatory molecules), route of administration, but also tumor factors, such as perfusion and the antigen expression levels and heterogeneity may impact viability and distribution characteristics and affect the interpretation of results.

In conclusion, each of the investigated approaches assesses distinct aspects of CD8^+^ T-cell distributions and infiltration of tumor lesions (e.g., number and distribution of residing CD8^+^ T cells or permissiveness for CD8^+^ T cell infiltration). We show that PET using anti-CD8 targeting tracers and *ex vivo* labeled cells can be used to assess the abundance of CD8^+^ T cells in the tumor, whereas all approaches appear suitable for evaluating activated lymph nodes. Our study underlines that selecting the optimal imaging approach depends on the research question and application of interest.

## Abbreviations

%ID/g: percentage injected dose per gram tissue
%ID: injected dose
RT: room temperature
Ax: container based on x%of maximum activity
AxLN: contralateral axial lymph node
IHC: immunohistochemistry
iv: intravenously
IL2: interleukin-2
InLN: inguinal lymph node
iTLC: instant thin-layer chromatography
LE: Labeling efficiency
LN: lymph nodes
MIP: maximum-intensity projections
PBS: phosphate buffered saline
PFE: PBS supplemented with 2%FCS and 1 mM EDTA
pi: post injection
SUV: standardized uptake values
TDLN: tumor draining lymph node
VOI: volume of interest

## Declarations

## Acknowledgments

We thank Daan Boreel, Annemarie Kip, Cathelijne Frielink, Bianca Lemmers-van de Weem, Iris Lamers-Elemans, and Kitty Lemmens-Hermans for technical assistance with the animal experiments. We thank Daphne Lobeek for assistance with the quantification of the PET data. We thank Paul Rijcken for assistance with the IHC image production.

## Authors’ contributions

Conceptualization: GGWS, RR, IA, JB, GA, MV, EA, SH. Data collection: GGWS, RR, IA, MB, LC, GF, JMK, PW, IH. Data analysis: GGWS, RR, IA, MB, LC, GF, JMK, PW, IH. Writing - original draft preparation: GGWS. Writing - review and editing: all authors.

## Funding

The research leading to these results received funding from the Innovative Medicines Initiatives 2 Joint Undertaking under grant agreement No 116106. This Joint Undertaking receives support from the European Union’s Horizon 2020 research and innovation programme and EFPIA. MV is recipient of ERC Starting grant CHEMCHECK (679921) and a Gravity Program Institute for Chemical Immunology tenure track grant by NWO. This communication reflects the author’s view and that neither IMI nor the European Union, EFPIA, or any Associated Partners are responsible for any use that may be made of the information contained therein.

## Competing interest

The authors have declared that no competing interest exists.

## Ethics approval

All *in vivo* experiments were approved by the Animal Welfare Body of the Radboud University, Nijmegen, and the Central Authority for Scientific Procedures on Animals (AVD1030020173885) and were performed in accordance with the principles stated by the Dutch Act on Animal Experiments (2014).

## Data availability statement

Data are available from the authors upon reasonable request.

## Notes

### Competing Interest Statement

The authors have declared no competing interest.

