## Supplemental material for "Head-to-head comparison of nuclear imaging approaches to quantify tumor CD8^+^ T-cell infiltration"

### Supplementals

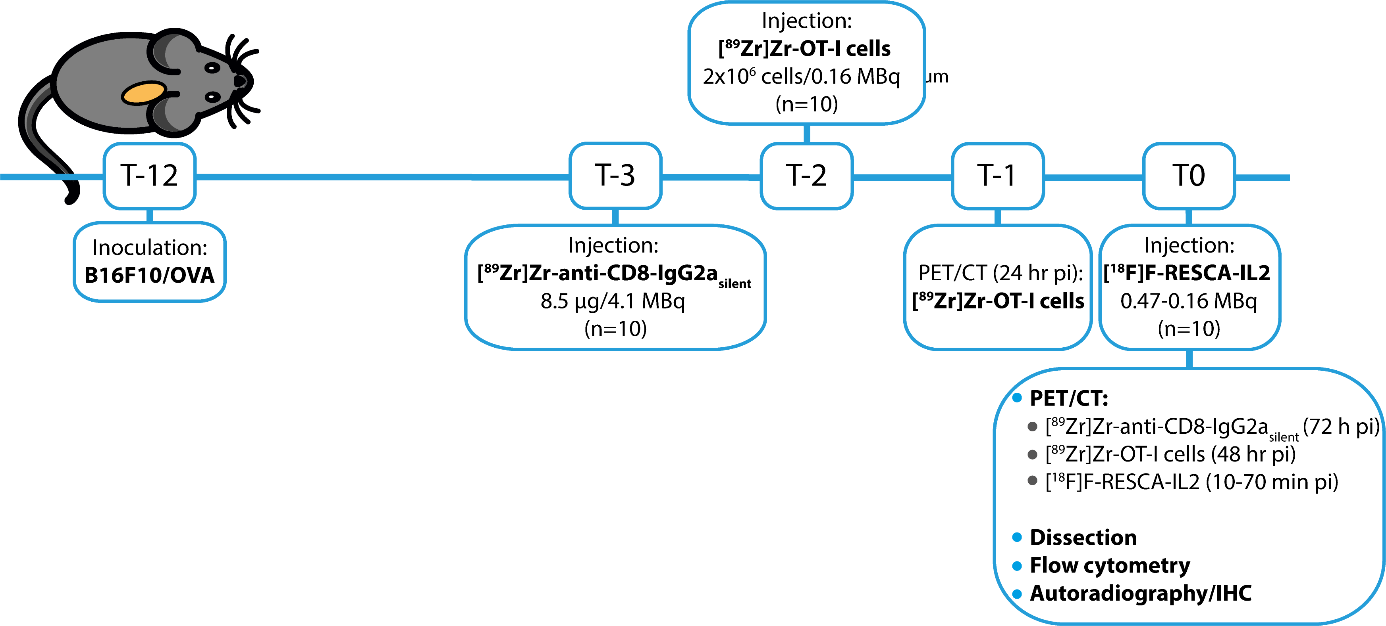
**Supplemental figure 1. Experimental time-line (in days).** C57BL/6 mice (n=36) were injected subcutaneously with B16F10/OVA cells. Three groups of mice (n=10/each) were injected with 8.5 µg [^89^Zr]Zr-anti-CD8-IgG2a_silent_, 2x10^6^ [^89^Zr]Zr-OT-I T cells, or [^18^F]AlF-RESCA-IL2, at t=-3, t=-2, or t=0, respectively. PET/CT imaging was performed at t=-1 for [^89^Zr]Zr-OT-I T cells injected mice and at t=0 for all mice. Subsequently, mice were euthanized followed by collection of relevant organs for *ex vivo* biodistribution analysis. One tumor halve was used for flow cytometric analysis of CD8^+^ T cell presence and one tumor halve was snap frozen for subsequent immunohistochemical and autoradiographic analysis.

**Supplemental table 1. Overview of biodistribution data for [^89^Zr]Zr-anti-CD8-IgG2a_silent_, [^89^Zr]Zr-OT-I CD8^+^ T cells, and [^18^F]AlF-RESCA-IL2.** *Ex vivo* biodistribution analysis in C57BL/6 mice bearing B16F10/OVA tumors injected with [^89^Zr]Zr-anti-CD8-IgG2a_silent_, from donor mice obtained and *ex vivo* labeled [^89^Zr]Zr-OT-I cells, or [^18^F]AlF-RESCA-IL2 at 72 h, 48 h, or 75 min post iv injection, respectively. Tissue uptake is presented in %ID/g. TDLN: tumor-draining, AxLN: axial, InLN: Inguinal lymph node, BAT: brown adipose tissue.

|  | **[^89^Zr]Zr-anti-CD8-IgG2a_silent_** | **[^89^Zr]Zr-OT-I CD8^+^ T cells** | **[^18^F]AlF-RESCA-IL2** |
| --- | --- | --- | --- |
|  | (%ID/g) | (%ID/g) | (%ID/g) |
| Blood | 1.5 ± 0.1 (n=10) | 0.4 ± 0.0 (n=9) | 2.8 ± 0.7 (n=10) |
| Muscle | 0.1 ± 0.1 (n=10) | 0.8 ± 0.5 (n=9) | 1.0 ± 0.2 (n=10) |
| Tumor | 3.4 ± 1.5 (n=10) | 1.9 ± 0.4 (n=9) | 1.7 ± 0.3 (n=10) |
| Spleen | 251.1 ± 26.5 (n=10) | 584.9 ± 53.0 (n=9) | 25.8 ± 4.6 (n=10) |
| TDLN | 123.4 ± 7.1 (n=5) | 171.9 ± 36.3 (n=9) | 2.5 ± 0.8 (n=9) |
| AxLN | 92.3 ± 26.0 (n=10) | 139.3 ± 30.5 (n=9) | 2.6 ± 1.3 (n=10) |
| InLN | 89.1 ± 36.4 (n=9) | 191.7 ± 44.6 (n=9) | 2.3 ± 0.4 (n=10) |
| Bone marrow | 6.0 ± 1.5 (n=10) | 48.6 ± 10.2 (n=9) | 6.2 ± 1.0 (n=10) |
| Thymus | 3.6 ± 0.7 (n=5) | 1.8 ± 0.3 (n=9) | 2.3 ± 0.4 (n=10) |
| Lung | 1.2 ± 0.2 (n=10) | 1.5 ± 0.4 (n=9) | 27.7 ± 4.0 (n=10) |
| Kidney | 2.2 ± 0.3 (n=10) | 2.7 ± 0.1 (n=9) | 115.0 ± 18.8 (n=10) |
| Liver | 6.9 ± 0.6 (n=10) | 41.5 ± 2.0 (n=9) | 35.6 ± 3.1 (n=10) |
| Duodenum | 8.4 ± 1.5 (n=10) | 1.1 ± 0.4 (n=9) | 14.1 ± 4.3 (n=10) |
| Colon | 1.5 ± 1.8 (n=10) | 0.8 ± 0.3 (n=9) | 2.6 ± 0.5 (n=10) |
| BAT | 0.5 ± 0.1 (n=10) | 1.8 ± 0.4 (n=9) | 2.9 ± 0.4 (n=10) |
| Femur | 2.2 ± 1.0 (n=10) | 13.6 ± 2.6 (n=9) | 4.4 ± 0.5 (n=10) |
| Knee | 2.1 ± 0.2 (n=10) | 14.4 ± 2.2 (n=9) | 5.3 ± 0.6 (n=10) |

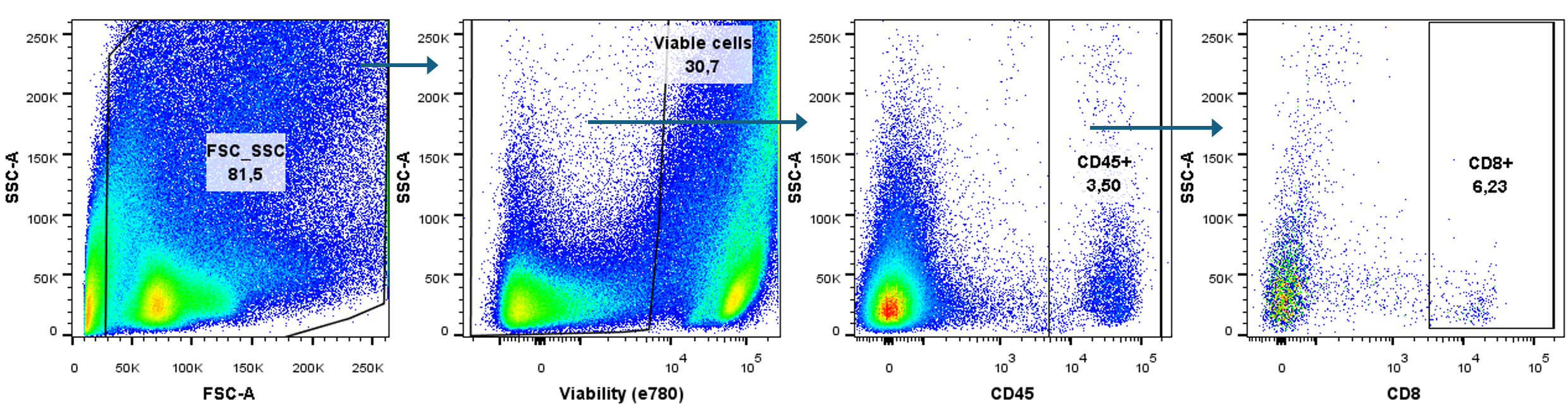
**Supplemental figure 2. Gating strategy for flow cytometric analysis of intratumoral CD8^+^ T cells.** Representative example of the gating strategy employed to evaluate the presence of T cell populations in B16F10/OVA tumors of C57BL/6 mice injected with either [^89^Zr]Zr-anti-CD8-IgG2a_silent_, from donor mice obtained and *ex vivo* labeled [^89^Zr]Zr-OT-I CD8^+^ T cells, or [^18^F]AlF-RESCA-IL2. Cells of interest were identified based on consecutive gating for 1) size and granularity, 2) viability, 3) expression of CD45, and 4) expression of CD8. SSC-A: Side Scatter parameter, FSC-A: Forward Scatter parameter.

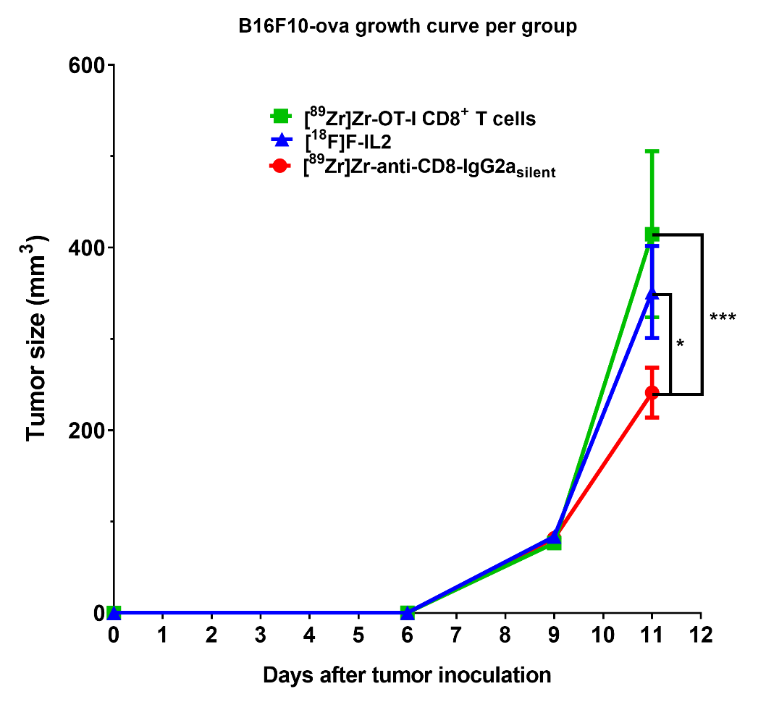

**Supplemental figure 3. Tumor sizes per treatment group over time.** Caliper measured volumes of B16F10-ova tumors (mm^3^) in C57BL/6 mice injected iv with [^89^Zr]Zr-anti-CD8-IgG2a_silent_, from donor mice obtained and *ex vivo* labeled [^89^Zr]Zr-OT-I CD8^+^ T cells, and [^18^F]ALF-RESCA-IL2 at day 9, 10, and 12, respectively. Data shown as mean±SEM.

**Supplemental table 2**. Overview of the tumor uptake in %ID/g from the biodistribution analysis juxtaposed with the calculated SUV values from data derived from the biodistribution analysis (SUV_biod_) and the PET quantification (SUV_5_, SUV_40_, SUV_max_). SUVs were calculated in both instances as a ratio between the activity per unit of volume in the tumor (activity/ml) and the activity per unit of whole body volume (Injected activity/gram of body weight). Tumor uptake values are shown (per mouse) for [^89^Zr]Zr-anti-CD8-IgG2a_silent_, from donor mice obtained and *ex vivo* labeled [^89^Zr]Zr-OT-I CD8^+^ T cells, and [^18^F]AlF-RESCA-IL2 in C57BL/6 mice bearing B16F10/OVA tumors. SUV: Standardized uptake value.

|  |  | **Biodistribution** | | | **PET quantification** | | |
| --- | --- | --- | --- | --- | --- | --- | --- |
|  | # | %ID/g | body weight (g) | SUV_biod_ | SUV_5_ | SUV_40_ | SUV_max_ |
| [^89^Zr]Zr-anti-CD8-IgG2a_silent_ | 1 | 2.187 | 22.4 | 0.490 | 0.356 | 0.732 | 0.881 |
|  | 2 | 3.851 | 21.2 | 0.816 | 0.508 | 0.766 | 1.216 |
|  | 3 | 3.534 | 20.9 | 0.739 | 0.386 | 0.714 | 1.003 |
|  | 4 | 7.171 | 21.2 | 1.520 | 0.555 | 0.939 | 1.706 |
|  | 5 | 2.211 | 19.2 | 0.425 | 0.381 | 0.700 | 0.935 |
|  | 6 | 4.930 | 20.7 | 1.021 | 0.606 | 0.911 | 1.547 |
|  | 7 | 2.427 | 20.7 | 0.502 | 0.355 | 0.740 | 0.965 |
|  | 8 | 2.808 | 20.7 | 0.581 | 0.436 | 0.769 | 0.954 |
|  | 9 | 2.852 | 20.5 | 0.585 | 0.461 | 0.785 | 0.903 |
|  | 10 | 2.414 | 19.4 | 0.468 | 0.364 | 0.651 | 0.732 |
| [^89^Zr]Zr-OT-I cells (48 hr) | 1 | 2.504 | 19.9 | 0.498 | 0.227 | 0.755 | 1.034 |
|  | 2 | 1.899 | 21.1 | 0.401 | 0.302 | 0.913 | 1.711 |
|  | 3 | 1.654 | 21.6 | 0.357 | 0.297 | 1.015 | 1.627 |
|  | 4 | 1.472 | 19.6 | 0.288 | 0.238 | 0.852 | 1.197 |
|  | 5 | 1.405 | 20.2 | 0.284 | 0.227 | 0.811 | 1.104 |
|  | 6 | 1.436 | 21.2 | 0.305 | 0.285 | 0.925 | 1.828 |
|  | 7 | 1.776 | 21.4 | 0.380 | 0.293 | 0.902 | 1.557 |
|  | 8 | 2.531 | 21.3 | 0.539 | 0.245 | 0.986 | 1.183 |
|  | 9 | 2.114 | 22 | 0.465 | 0.214 | 0.839 | 0.885 |
| [^18^F]AlF-RESCA-IL2 | 1 | 1.426 | 21.7 | 0.310 | 0.166 | 0.548 | 0.642 |
|  | 2 | 2.211 | 20.4 | 0.451 | 0.244 | 0.583 | 1.008 |
|  | 3 | 1.916 | 21.1 | 0.404 | 0.291 | 0.738 | 1.435 |
|  | 4 | 2.089 | 21.2 | 0.443 | 0.224 | 0.573 | 0.862 |
|  | 5 | 1.048 | 21.8 | 0.228 | 0.155 | 0.611 | 0.723 |
|  | 6 | 1.555 | 22.3 | 0.347 | 0.213 | 0.704 | 1.279 |
|  | 7 | 1.810 | 21 | 0.380 | 0.190 | 0.573 | 0.689 |
|  | 8 | 1.652 | 19.5 | 0.322 | 0.180 | 0.550 | 0.715 |

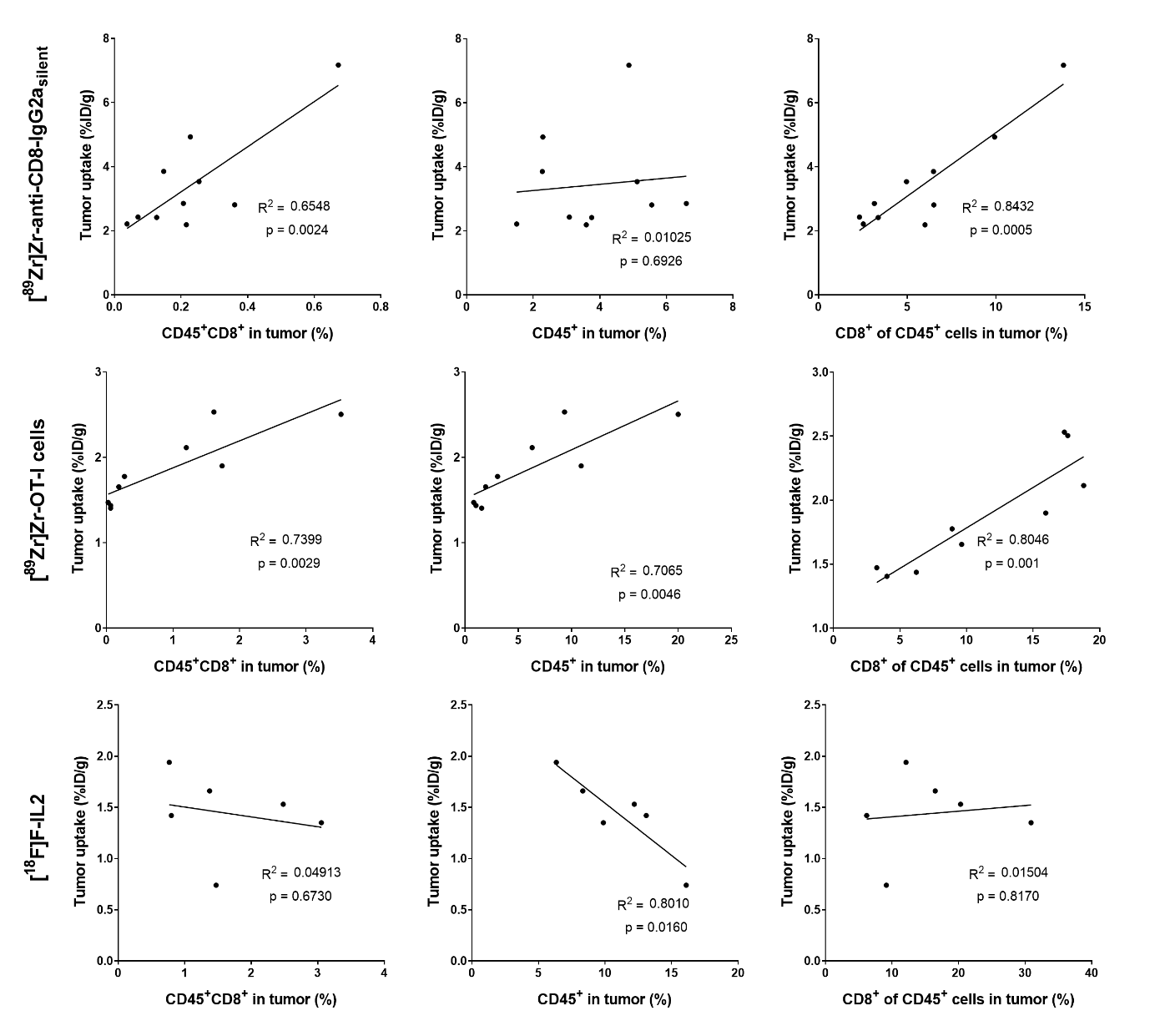
**Supplemental figure 4. T cell populations in the tumor versus tumor uptake per tracer.** Pearson’s correlation analysis of the presence of T cell populations and tumor uptake of [^89^Zr]Zr-anti-CD8-IgG2a_silent_, from donor mice obtained and *ex vivo* labeled [^89^Zr]Zr-OT-I CD8^+^ T cells, and [^18^F]AlF-RESCA-IL2 in C57BL/6 mice bearing B16F10/OVA tumors. Tumor uptake (%ID/g) and the presence of T cell populations was determined by *ex vivo* biodistribution analyses and flow cytometry, respectively. Shown are the correlations of tumor uptake with: 1) %CD45+CD8+ cells of the viable population, 2) %CD45^+^ of viable cells, and 3) %CD8^+^ of CD45^+^ cells within the viable cell population.

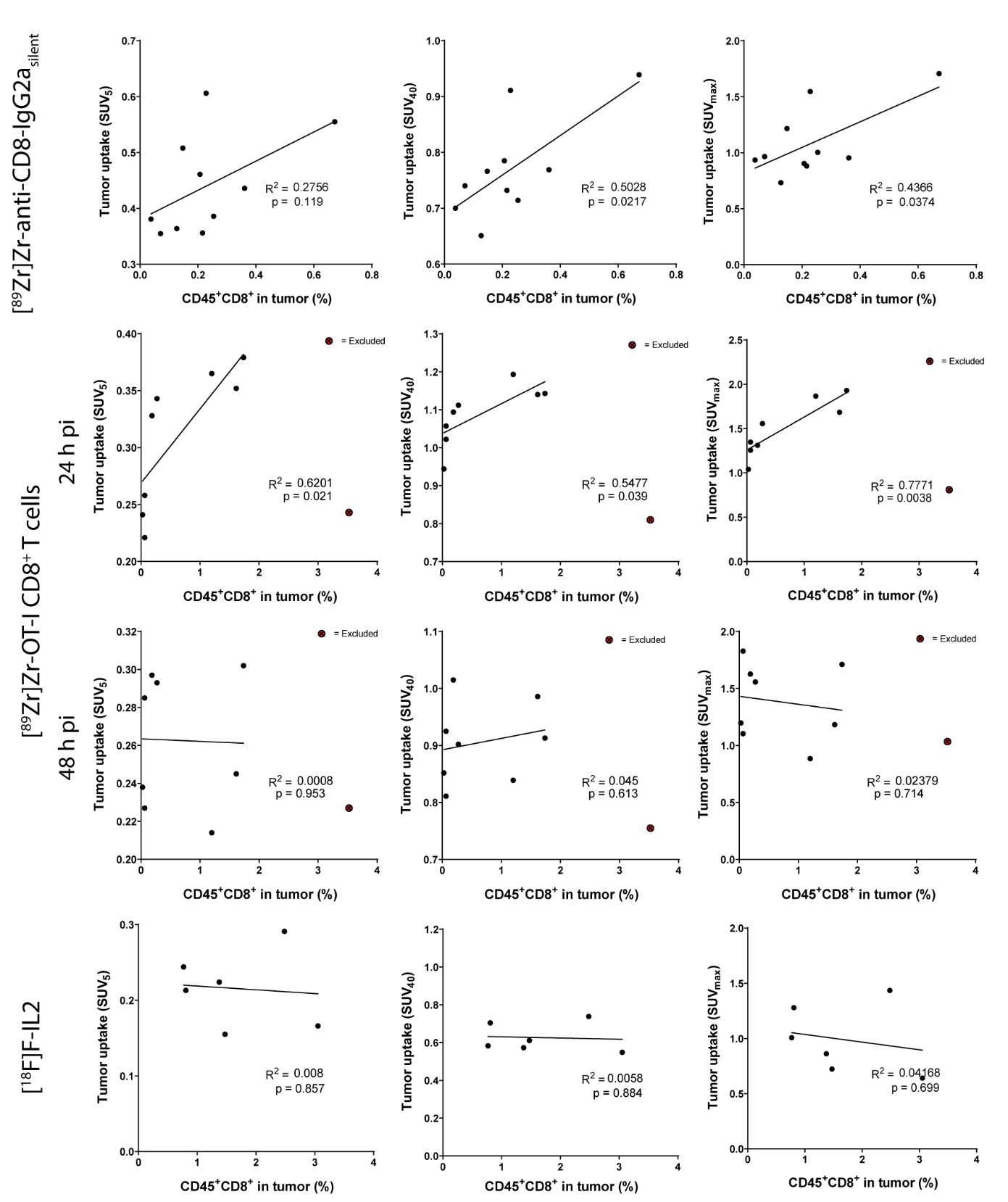

**Supplemental figure 5. Correlation analyses of the number of tumor infiltrated CD8^+^ T cells with the PET quantified tumor uptake of [^89^Zr]Zr-anti-CD8-IgG2a_silent_ antibody, [^89^Zr]Zr-OT-I CD8^+^ T cells, and [^18^F]AlF-RESCA-IL2.** Pearson’s correlation analysis of the presence of CD45^+^CD8^+^ T cells and tumor uptake of [^89^Zr]Zr-anti-CD8-IgG2a_silent_, from donor mice obtained and *ex vivo* labeled [^89^Zr]Zr-OT-I CD8^+^ T cells, and [^18^F]AlF-RESCA-IL2 in C57BL/6 mice bearing B16F10/OVA tumors. The correlation of the percentage of CD45^+^CD8^+^ T cells as determined by flow cytometry with their tumor uptake (SUV_5_, SUV_40_, and SUV_max_) determined by PET image quantification.

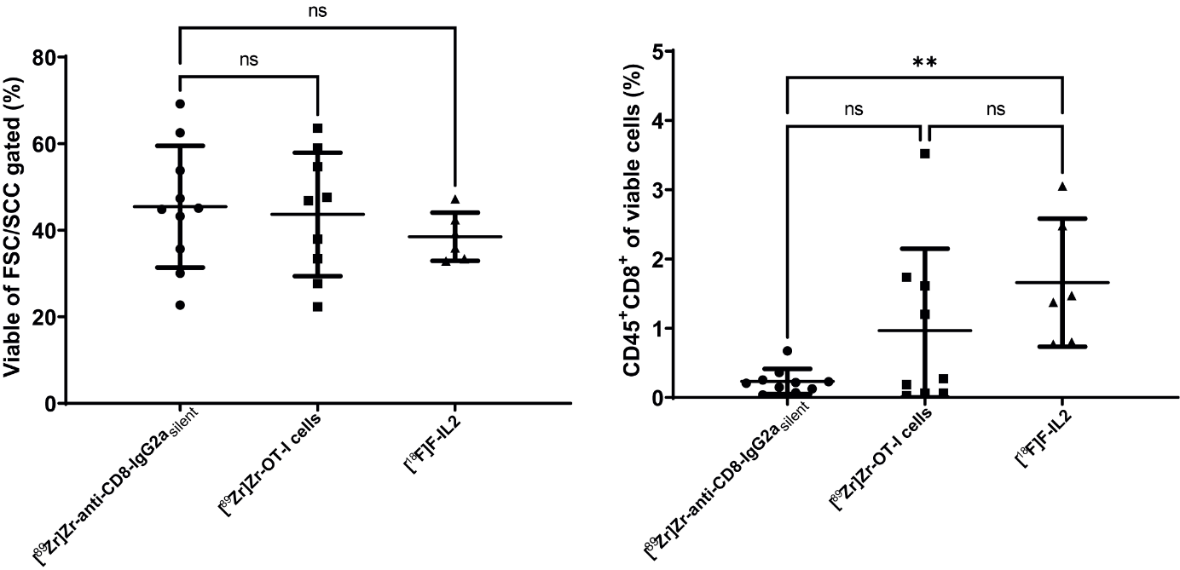
**Supplemental figure 6. Cell populations in the tumor.** The viability of obtained cells and the presence of CD45^+^CD8^+^ T cells in B16F10/OVA tumors of C57BL/6 mice injected with either [^89^Zr]Zr-anti-CD8-IgG2a_silent_, from donor mice obtained and *ex vivo* labeled [^89^Zr]Zr-OT-I CD8^+^ T cells, or [^18^F]AlF-RESCA-IL2. Depicted are, as determined by flow cytometry, the percentage of viable cells within the total cell population (left panel) and percentage CD45^+^CD8^+^ T cells within the total viable population (right panel).

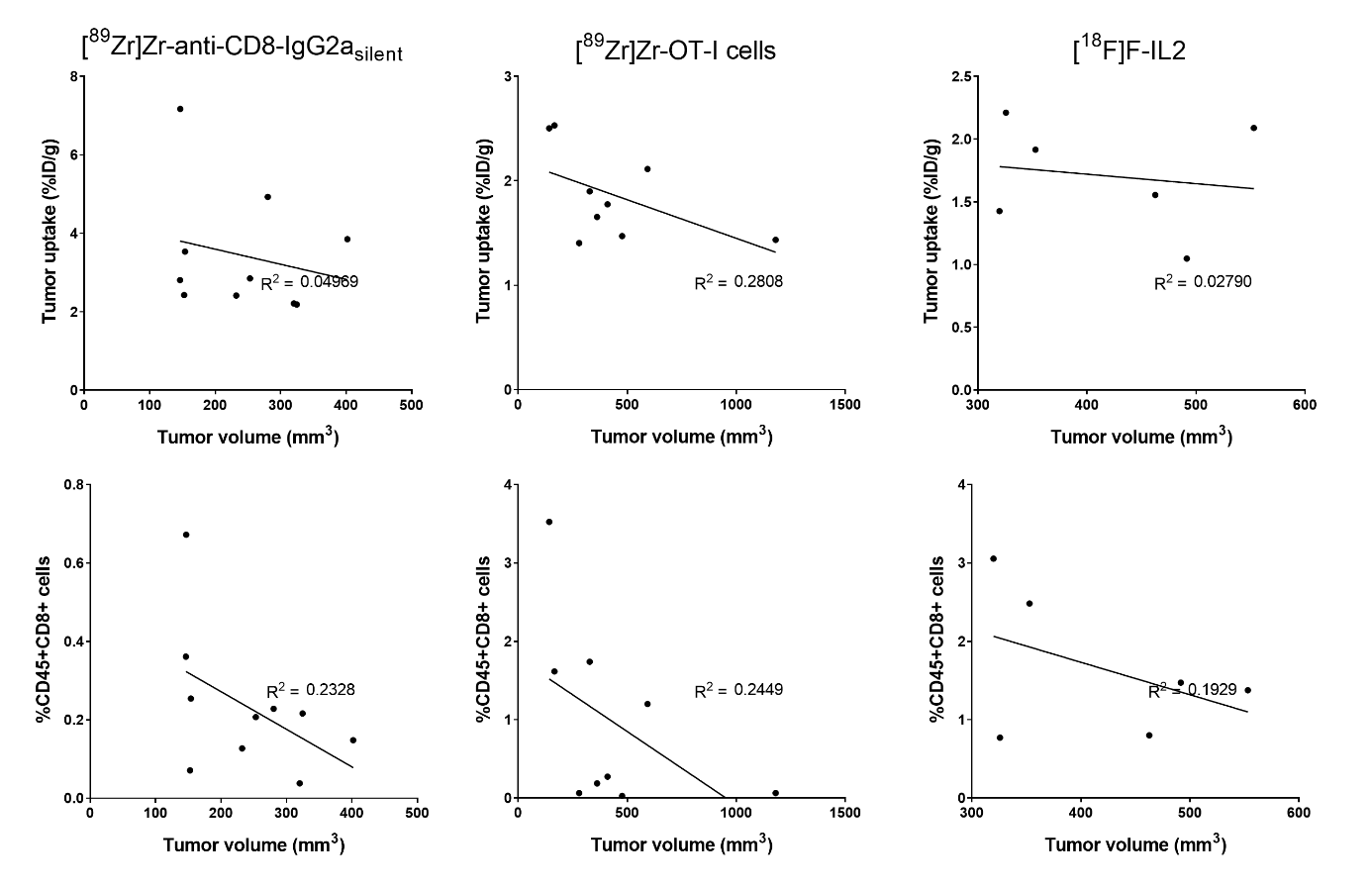
**Supplemental figure 7. Correlation analysis of tumor volume with the tumor uptake and fraction of tumor residing CD8^+^ T cells.** Pearson’s correlation analysis of the tumor volume with: 1) tumor uptake of [^89^Zr]Zr-anti-CD8-IgG2a_silent_, from donor mice obtained and *ex vivo* labeled [^89^Zr]Zr-OT-I CD8^+^ T cells, and [^18^F]AlF-RESCA-IL2, and 2) the %CD45^+^CD8^+^ T cells in C57BL/6 mice bearing B16F10/OVA tumors. Tumor volume, tumor uptake (%ID/g), and the %CD45^+^CD8^+^ T cells was determined by caliper measurements, *ex vivo* biodistribution analyses, and flow cytometry, respectively.

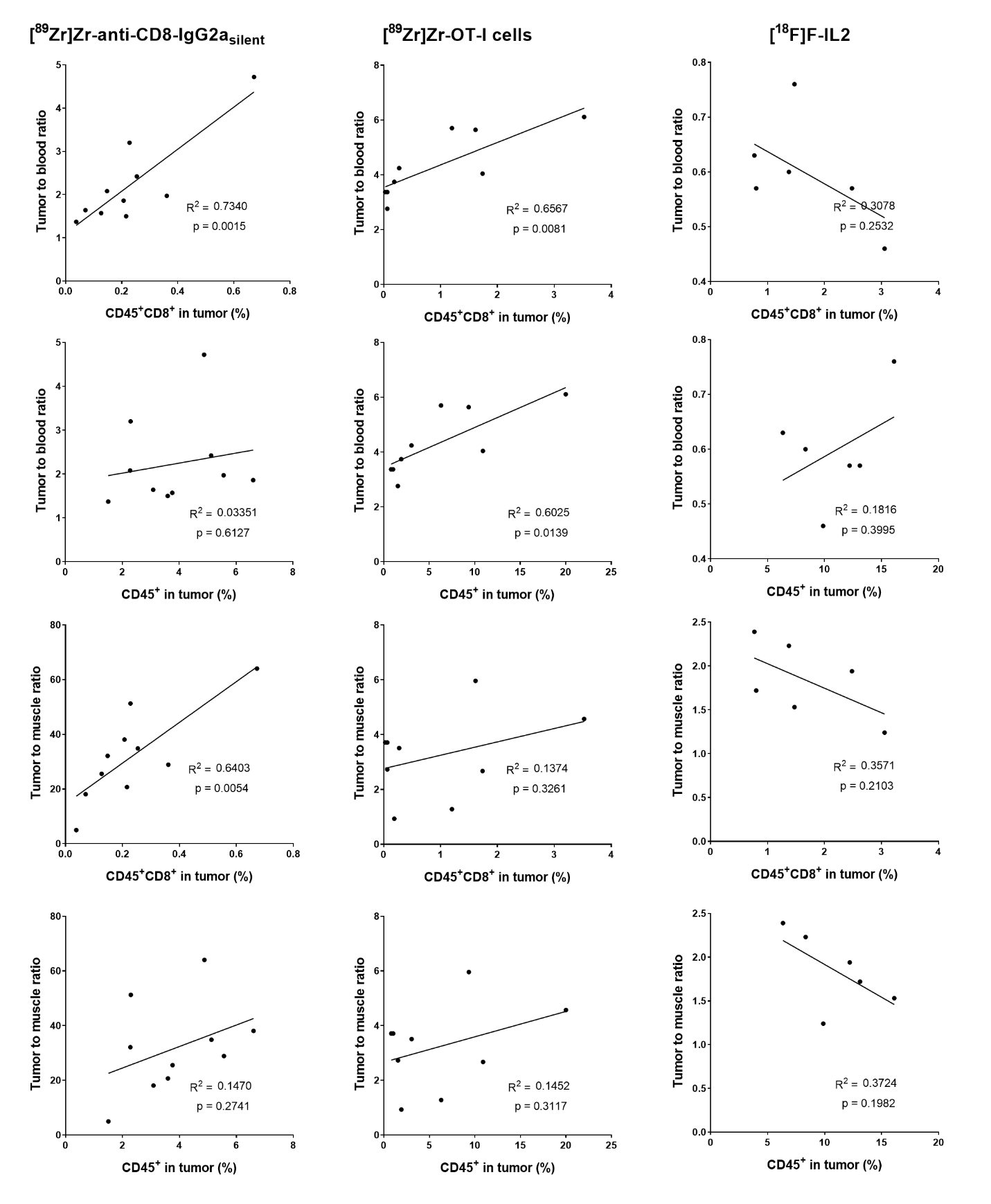

**Supplemental figure 8. Correlation analyses of the number of tumor infiltrated T cells with the tumor-to-blood and tumor-to-muscle ratios of [^89^Zr]Zr-anti-CD8-IgG2a_silent_ antibody, [^89^Zr]Zr-OT-I CD8^+^ T cells, and [^18^F]ALF-RESCA-IL2.** Pearson’s correlation analysis of the presence of CD45^+^CD8^+^ and CD45^+^ T cells and with biodistribution analysis evaluated tumor-to-blood and tumor-to-muscle ratios of [^89^Zr]Zr-anti-CD8-IgG2a_silent_, from donor mice obtained and *ex vivo* labeled [^89^Zr]Zr-OT-I CD8^+^ T cells, and [^18^F]AlF-RESCA-IL2 in C57BL/6 mice bearing B16F10/OVA tumors.
